# Structural basis of Nipah virus replication

**DOI:** 10.1101/2024.10.04.616610

**Authors:** Fernanda A. Sala, Katja Ditter, Olexandr Dybkov, Henning Urlaub, Hauke S. Hillen

**Affiliations:** University Medical Center Göttingen, Department of Cellular Biochemistry, Humboldtallee 23, D-37073 Göttingen, Germany; Max Planck Institute for Multidisciplinary Sciences, Research Group Structure and Function of Molecular Machines, Am Fassberg 11, D-37077 Göttingen, Germany; Max Planck Institute for Multidisciplinary Sciences, Bioanalytical Mass Spectrometry Group, Am Fassberg 11, D-37077, Göttingen, Germany; University Medical Center Göttingen, Institute for Clinical Chemistry, Bioanalytics Group, Robert-Koch-Straße 40, D-37077 Göttingen, Germany; Cluster of Excellence “Multiscale Bioimaging: from Molecular Machines to Networks of Excitable Cells” (MBExC), University of Göttingen, D-37075 Göttingen, Germany; Göttingen Center for Molecular Biosciences (GZMB), Research Group Structure and Function of Molecular Machines, University of Göttingen, D-37077 Göttingen, Germany

## Abstract

Nipah virus (NiV) is a non-segmented negative-strand RNA virus (nsNSV) with high pandemic potential, as it frequently causes zoonotic outbreaks and can be transmitted from human to human. Its RNA-dependent RNA polymerase (RdRp) complex carries out viral genome replication and transcription and is therefore an attractive drug target. However, to date no structural data is available on the NiV RdRp complex. Here, we report cryo-EM structures of NiV RdRp in the apo and in an early elongation state with RNA and incoming substrate bound. The structure of the apo enzyme reveals the architecture of the NiV RdRp complex, which shows a high degree of similarity to other nsNSV RdRps. The structure of the RNA-bound NiV RdRp shows how the enzyme interacts with template and product RNA during early replication and how nucleoside triphosphates are bound in the active site. Comparisons show that RNA binding leads to rearrangements of key elements in the RdRp core and to ordering of the flexible C-terminal domains of NiV L required for RNA capping. Taken together, these results reveal the first structural snapshots of an actively replicating nsNSV RdRp and provide insights into the mechanisms of genome replication and transcription by NiV and related viruses.

## INTRODUCTION

Nipah virus (NiV) is an emerging zoonotic pathogen of the Paramyxoviridae family within the order Mononegavirales, which encompasses nonsegmented, negative-strand RNA viruses (nsNSVs). Members of this order also include other important human pathogens such as respiratory syncytial virus (RSV), vesicular stomatitis virus (VSV), rabies virus (RABV) and Ebola virus (EBOV)^1^. Since its initial identification in the 1990s in Malaysia and Singapore^2^, NiV has caused recurrent outbreaks, the most recent in Bangladesh and India. The virus represents a major public health concern due to its high mortality rates (up to 80 %) and human-to-human transmission^3^. Currently, no vaccines or targeted antiviral treatments to combat Nipah virus infection and disease are available. As a result, the World Health Organization (WHO) has classified Nipah virus as a priority for epidemic preparedness as part of its Blueprint for Action to Prevent Epidemics^4^.

Central to the life cycle of NiV is its RNA-dependent RNA polymerase (RdRp) complex. As in other nsNSVs, it is comprised of the L protein, a large multi-functional polypeptide that contains all required enzymatic activities, and its cofactor, the P protein (called VP35 in Filoviridae)^5^. This machinery carries out both genome replication as well as transcription of individual genes. In both cases, the RdRp initiates RNA synthesis *de novo* at a leader sequence (*Le*) at the 3’ end of the (-)-sense genome. For transcription, the RdRp then undergoes reinitiation at the start of each of the six genes and produces capped and poly-adenylated mRNAs. By contrast, during replication, the RdRp produces a fulllength (+)-sense antigenome, which in turn serves as template for synthesis of a (-) sense genome copy^6^. Due to its central role for both transcription and replication of the Nipah genome, the RdRp complex represents an attractive target for the development of antiviral compounds.

The L protein is comprised of three enzymatic and two structural domains: the RNA-dependent RNA polymerase (RdRp) domain, responsible for catalyzing RNA synthesis, the polyribonucleotidyltransferase (PRNTase or CAP) domain involved in cap addition, the connector domain (CD), the methyltransferase (MTase) domain crucial for cap methylation, and the C-terminal domain (CTD)^7^. The P protein consists of an N-terminal domain (NTD), a central oligomerization domain (OD), and a C-terminal X domain (XD). While a crystal structure of the oligomerization domain of the NiV P protein has been reported^8^, no structural data on the intact NiV RdRp complex is available.

Over the past years, structures of RdRp complexes from several members of the Paramyxoviridae family as well as other nsNSVs have been determined in the apo state without nucleic acids bound^9–16^. In addition, structures of EBOV and RSV RdRps in complex with leader RNA have recently been obtained^17,18^. These structures have shed light on the overall architecture of nsNSV RdRps and provided first insights into how they interact with the template RNA during replication initiation. However, to date, no structures of actively replicating or transcribing nsNSV RdRps have been reported. As a consequence, the molecular mechanisms underlying nsNSV genome replication and transcription and how these different functions are regulated remain poorly understood.

Here, we present single-particle cryo-electron microscopy (cryo-EM) structures of the NiV RdRp complex in its apo state as well as in an early elongating state, with template RNA, product and incoming nucleotide triphosphate (NTP) bound. These structures provide the first molecular insights into Nipah virus replication and represent the first structures of an nsNSV in the actively replicating state. Comparison between the two structures reveals key functional elements that undergo conformational rearrangements upon RNA binding and early elongation, and show that changes cause large-scale structural reorganization of the RdRp complex, including ordering of the otherwise flexible C-terminal domains. Taken together, these results provide a framework for understanding and targeting Nipah virus replication and transcription and represent an important milestone in our understanding of reaction cycle of nsNSV RdRps in general.

## RESULTS

### Structure of the apo NiV RdRp complex

To obtain structural insights on the NiV RdRp complex, we co-expressed codon-optimized NiV L and P proteins in insect cells. Purification via an affinity tag on the NiV L protein led to co-purification of NiV P protein, and we further purified the complex to homogeneity by heparin and size exclusion chromatography and confirmed its identify using mass-spectrometry (**Supplementary Figure 1a,b, Methods**). Incubation of the complex with a synthetic single-stranded RNA comprising the NiV 3’ leader sequence (*Le*) and substrate NTPs led to the formation of RNA products of the expected size range, showing that the purified NiV RdRp is active (**Supplementary Figure 1c**). We then analyzed the RdRp complex by single-particle cryo-EM, which led to a reconstruction at an overall resolution of 2.6 Å (**Supplementary Figure 2, Supplementary Table 1**). This allowed us to build and refine an atomic model of the apo NiV RdRp complex with excellent stereochemistry (**Figure 1, Supplementary Table 1**).

**Fig. 1.**
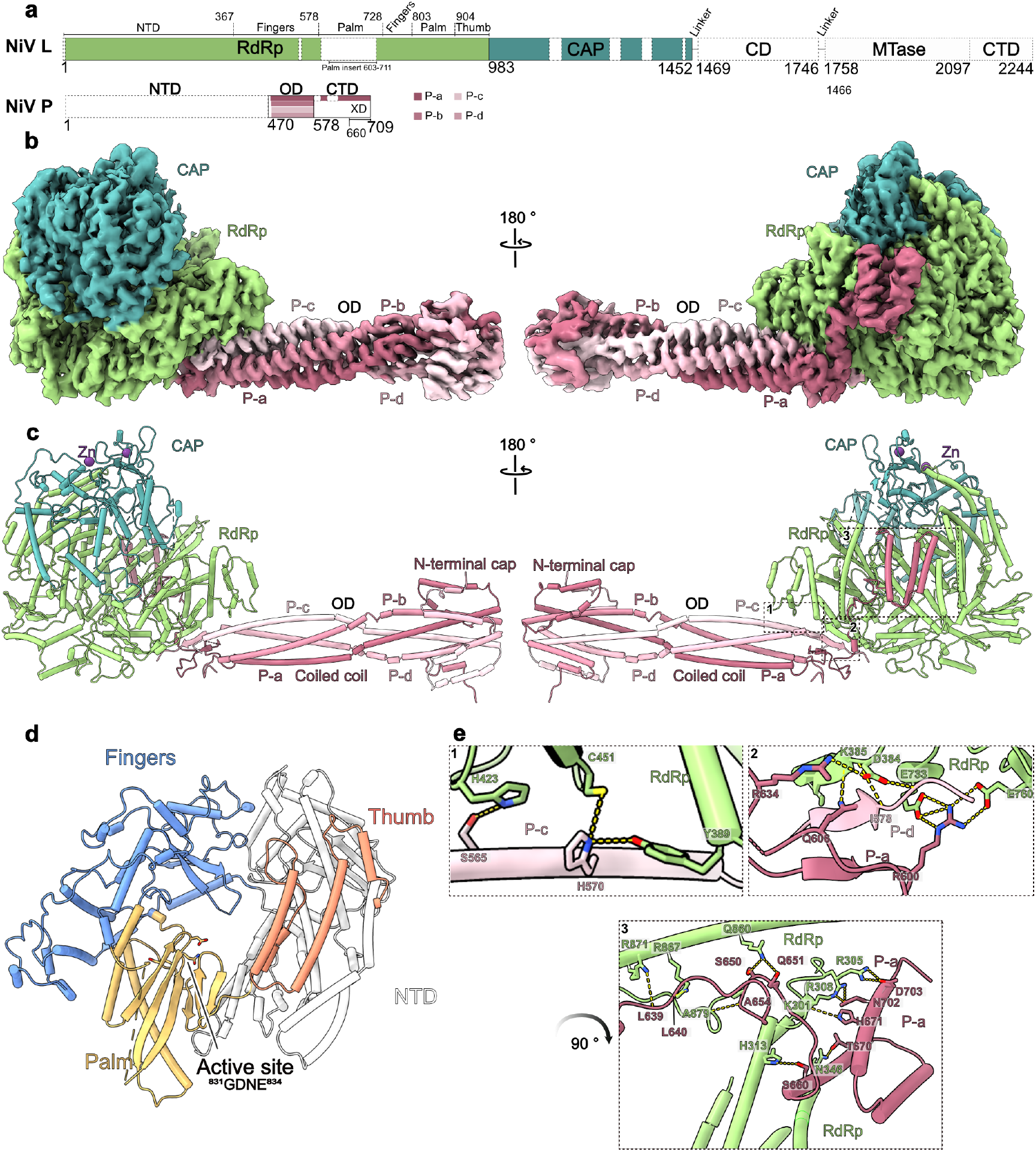
Structure of the apo NiV RdRp complex. **a)** Schematic domain representation of NiV L and P protein. The RdRp domain of NiV L is shown in green and the CAP domain in dark cyan. The four P proteins are shown in varying shades of pink. Unmodelled regions are indicated as white dashed boxes. Numbers give amino acid positions at domains boundaries. This color scheme is consistently applied throughout. **b)** Cryo-EM density map of the apo-NiV RdRp complex (Local resolution filtered map). **c)** Cartoon representation of the apo NiV RdRp complex with Zn^2+^ ions shown as purple spheres. **d)** Structure of the NiV L RdRp domain colored by subdomains. The fingers domain is blue, the thumb is salmon, the palm is yellow gold, and the NTD is gray. The active site, represented by the GDNE motif (residues 831-834), is shown as sticks. **e)** Close-up view of interactions between the NiV P and L protein. The interactions are categorized into three regions (1-3) indicated in c. Interacting side chains are shown as sticks, with hydrogen bonds highlighted by yellow dashed lines. 1) Interaction between P-c an L RdRp. 2) Interactions among P-a, P-d, and L proteins. 3) Interaction between P-a XD and L RdRp.

The overall structure of the NiV RdRp complex resembles that previously observed for other nsNSV RdRps^5,6^. It is comprised of one copy of NiV L and four copies of NiV P (**Figure 1a,b,c**), a stochiometry that was previously observed for RdRp complexes of other paramyxoviruses^14,15,19^, rhabdoviruses^9,20^, pneumoviruses^12,13,16^ and filoviruses^11^. We observe clear density for the RdRp and CAP domains of NiV L (residues 1-1452), which together adopt a globular fold with a large cavity at its center. By contrast, the C-terminal CD, MTase and CTD of L are not visible, indicating that these domains are conformationally flexible in the apo state of the enzyme (**Figure 1a**). This is consistent with structural studies on some other nsNSV RdRp complexes, which also did not observe density for the corresponding parts of L^11–13,21^. For the NiV P protein, we observe density for the OD domains (residues 470-578) of all four copies in the complex, which assemble into a rod-like structure that protrudes from the RdRp core. For one of the NiV P protomers (P-a), we observe additional density for a linker and the C-terminal domain X domain (XD), which binds at the side of the NiV L RdRp domain. The remaining parts of NiV P are not visible in our maps, indicating conformational flexibility. This is consistent with structures of related nsNSV RdRp complexes, in which large parts of P were also not visible^9,12–16,19,20^.

The RdRp domain of NiV L adopts the canonical, highly-conserved right hand fold characteristic of single-subunit RNA polymerases, composed of thumb, fingers and palm subdomains (**Figure 1d**). The active site is located within a large cavity at the center of the RdRp domain and contains seven structural motifs (A-G) conserved across viral RdRps (**Supplementary Figure 3a**)^22^. The catalytic GDNE motif (res. 831-834) resides within motif C, which is part of the palm subdomain (**Figure 1d and Supplementary Figure 3a**). Additionally, the RdRp domain contains two conserved elements, the supporting helix (res. 588-600) and the supporting loop (res. 579-587), which have been observed to adopt different conformations associated with distinct functional states in other nsNSV RdRps^13,18,23^. While the supporting loop is partially ordered in the apo NiV RdRp complex, the supporting helix appears to be flexible (**Supplementary Figure 3b**). Compared to other nsNSV RdRps, NiV L contains a large sequence insertion of unknown function in its palm subdomain (res. 603-711) (**Supplementary Figure 4**) for which we do not observe density, indicating that it is conformationally flexible. The active site is accessible to the solvent through four channels, which presumably serve as NTP entry, template entry, template exit and product exit sites, respectively (**Supplementary Figure 3c**). The CAP domain, which contains the PRNTase activity required for RNA capping, resides on top of the RdRp domain and forms a lid over the active site cavity (**Figure 1c**). It contains two zinc-binding motifs (**Supplementary Figure 4**), which have been proposed to serve a primarily structural role in the related VSV RdRp complex^10^. It further contains two conserved structural elements: the priming loop (res. 1266-1289) and the intrusion loop (res. 1337-1362). The priming loop may be involved in initiation by interacting with the first incoming nucleotides^24,25^. The intrusion loop contains a catalytic HR motif, which forms a covalent bond with the 5’ end of the nascent RNA during capping^26,27^. Both elements have been observed to be either disordered or to adopt different conformations in apo structures of other viral RdRps (**Supplementary Figure 3d**)^9–11,14,15,20,23^. In the apo NiV RdRp complex, both the priming and intrusion loop display only weak density, indicating that they are largely disordered. Thus, neither of these elements appear to be ordered in the absence of nucleic acids.

The four protomers of NiV P form a characteristic stalk that extends from the polymerase core and is formed by coiled-coil interactions between their OD domains (**Figure 1b,c**), reminiscent of the structure of P/VP35 in other nsNSV RdRps^6^. A unique feature of the NiV P is a mushroom-like tip at the distal end of the stalk. Although this region is not well resolved in our reconstruction, we could model it with the help of the crystal structure of the NiV P OD domain tetramer^8^. This shows that the cap is formed by res. 478-506 of NiV P, which form helices that fold back onto the coiled-coil OD bundle (**Figure 1c, Supplementary Figure 3e**). The base of the P-stalk is anchored to the polymerase core mainly through interactions between the RdRp domain of NiV L and P-c and P-d, respectively (**Figure 1e**). The XD domain of P-a folds into a three-helix bundle that clamps onto the side of the NiV L RdRp domain, near the substrate NTP entry channel (**Figure 1e**). This is consistent with previous structures of EBOV^11^ and hPIV5^15^ apo RdRp complexes, in which the XD domains of VP35 or P were observed to bind in similar positions.

Overall, the structure of the apo NiV RdRp complex shows that it adopts a highly conserved architecture resembling that of other nsNSV RdRps and that its C-terminal domains as well as key functional elements in the RdRp core are mobile in the absence of nucleic acids.

### Structure of an actively replicating NiV RdRp complex

In order to obtain more detailed insights into the structural basis of Nipah virus replication, we next aimed to capture a structural snapshot of the NiV RdRp complex during active RNA synthesis. To achieve this, we incubated the recombinant NiV RdRp complex with a synthetic RNA comprising the NiV *Le* sequence in the presence of ATP, CTP, UTP and the ß,y non-hydrolyzable GTP analog Guanosine-5’-[(β,γ)-imido]triphosphate (GMPPNP). Single-particle cryo-EM analysis of this sample revealed two major particle populations **(Supplementary Figure 5)**. The first resembles the above-described apo NiV RdRp, with no RNA visible in the active site and disordered OD, MTase and CTD domains. The second shows clear extra density for a duplex RNA in the active site, thus representing an RNA-bound NiV RdRp complex. Refinement of this particle population led to a consensus reconstruction at an overall resolution of 2.8 Å. Multibody refinement in Relion led to improved local maps for the RdRp core, the P-stalk and the C-terminal domains, which enabled us to build a molecular model of RNA-bound NiV RdRp (**Supplementary Table 1**).

The core of the RNA-bound NiV RdRp complex adopts an identical overall structure as the apo enzyme (r.m.s.d 0.971 Å) (**Figure 2a,b,c, Supplementary Figure 6a**). Compared to the apo NiV RdRp, the reconstruction showed improved density for Niv P. First, Multibody refinement with a mask around the P-stalk led to an improved map for the mushroom-like tip of the stalk (**Supplementary Figure 6b**). Second, we were able to model a segment C-terminal of the OD domain of P-c, which meanders along the side of the NiV L RdRp domain (**Supplementary Figure 6c**). The P stalk adopts a slightly different orientation with regards to NiV L than in the apo structure, suggesting that it is conformationally dynamic (**Supplementary Figure 6d**). Most strikingly, in contrast to the apo NiV RdRp structure, we observe clear density for the previously disordered flexible domain of NiV L comprising the OD, MTase and CTD (**Figure 2a,b,c**). Thus, ordering of these domains appears to correlate with RNA binding. In the active site, characterized by the conserved right-hand fold domain containing the catalytic motifs (A-G), we observe density for a 9-base pair RNA duplex composed of the NiV *Le* template as well as product RNA (**Figure 2d,e**). The resolution is sufficient to unambiguously discriminate between purine and pyrimidine bases, revealing that the NiV RdRp has synthesized RNA up to the first occurrence of a cytidine base in the template strand (C10, numbering from 3’ to 5’) *in situ*. In addition, we observe density for three additional downstream nucleotides of the template RNA (C10-C12). The 3’ end of the nascent RNA is positioned in the -1 site and the + site is occupied by an incoming NTP, which base pairs with the templating cytidine base (C10) and which we modeled as GMPPNP (**Figure 2d**). We observe clear density for the sugar, base and for all three phosphates of GMPPNP, but no connection to the 3’ end of the nascent RNA is visible at reasonable map threshold, suggesting that we captured the incoming substrate prior to phosphodiester bond formation (**Figure 2d**). Thus, the structure represents an actively replicating NiV RdRp complex during early RNA elongation, captured in the post-translocated state with incoming NTP bound. This state has, to our knowledge, not been described for any nsNSV RdRp so far.

**Fig. 2.**
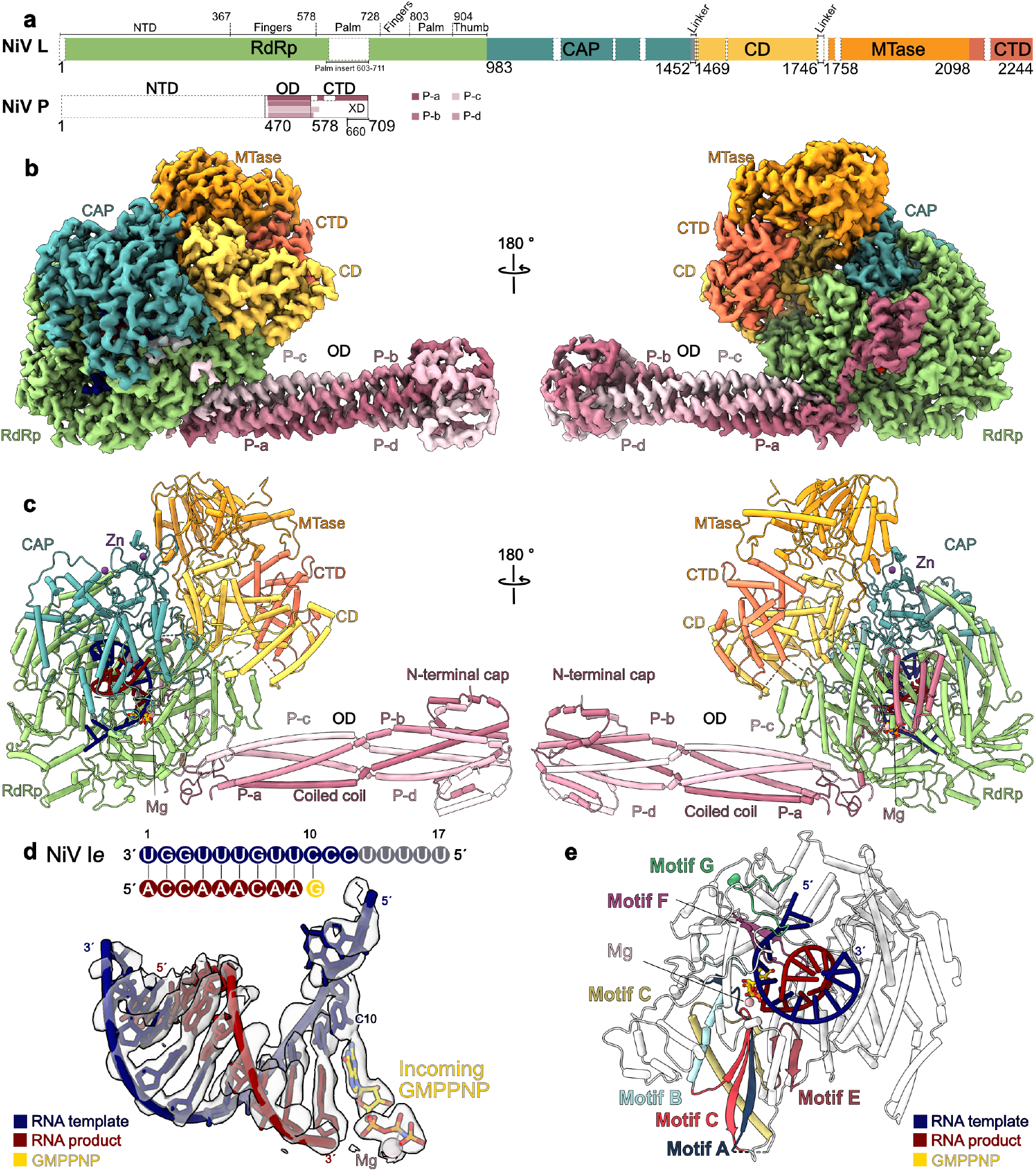
Structure of the actively replicating NiV RdRp complex. **a)** Schematic domain representation of NiV L and P protein. The RdRp domain of NiV L is shown in green, the CAP domain in dark cyan, CD in yellow, MTase in gold yellow and CTD in orange. The four P proteins are shown in varying shades of pink. Unmodelled regions are indicated as white dashed boxes. This color scheme is consistently applied throughout. **b)** Cryo-EM density maps of the actively replicating NiV RdRp complex (post-processed map). **c)** Cartoon representation of the actively replicating NiV RdRp complex. The template RNA is shown in blue and the product RNA in red. Zn^2+^ and Mg^2+^ are shown as purple and pink spheres in, respectively; Colour code used throughout. **d)** Schematic and stick representation of the RNA template and product with incoming GMPPNP (yellow) and corresponding cryo-EM density. **e)** Structure of the NiV L RdRp domain with RNA bound with conserved catalytic motifs color-coded from A to G. The active site GDNE motif (residues 831-834) and incoming GMPPNP are shown as sticks.

### Interactions between NiV RdRp and RNA during RNA synthesis

The structure of the actively replicating NiV RdRp complex reveals how the enzyme interacts with nucleic acids during early RNA synthesis. The template RNA enters the active site through the template entry channel, which is located at the interface between the RdRp and CAP domains (**Supplementary Figure 6e**). Comparison between the apo and replicating NiV RdRp structures shows that in the RNA-bound state, the fingers domain undergoes a shift of roughly 4 Å, which may stabilize the entering template RNA (**Supplementary Figure 6a**). The template strand follows the same trajectory as observed in the EBOV and RSV initiation complexes^17,18^, suggesting that no rearrangements of the template occur between initiation and the early stages of RNA synthesis (**Supplementary Figure 6f**).

The active site accommodates a 9-base pair (bp) RNA duplex, which is stabilized by interactions with residues from the palm, finger, thumb and CAP domain (**Figure 3a,b**). The upstream edge of the duplex faces towards a wall formed by res. 1050-1059 and 1117-1125, which likely limits the length of duplex that can be accommodated. The 5’ end of the product RNA faces towards the RNA exit channel, which is formed by the between the CAP and RdRp domains (**Supplementary Figure 6e**). In the active center of the enzyme, the templating base C10 is stabilized in the +1 position by stacking against the conserved residue F553 in motif F of the fingers domain (**Figure 3a, Supplementary Figure 4**). The incoming GMPPNP is stabilized through base pairing with the templating C10. In addition, it stacks against R551 from motif F, which protrudes towards the NTP binding site and is also strictly conserved in nsNSV RdRps (**Figure 3a, Supplementary Figure 4**). The triphosphate moiety of the incoming NTP binds in a groove formed by the catalytic loop in motif C (residues 826-837), which contains one of the two conserved catalytic aspartate residues (D832), and motif A, which contains a second conserved catalytic aspartate (D722). Both aspartates face towards the triphosphate and complex one of the two catalytic Mg ions, for which we observe density in between the aspartates and the triphosphate moiety (**Figure 2d**). Comparison between the apo and RNA-bound states of NiV RdRp shows that motif C repositions to accommodate the product RNA (**Figure 3c**). On the opposing side, the triphosphate of the incoming NTP is stabilized by interactions with R551. Structural comparison to other viral RdRps for which elongating structures have been reported, such as SARS-CoV-2^28^ and Influenza A^29^, shows that the architecture of the active site and arrangement of catalytic residues is highly conserved (**Supplementary Figure 6g**).

**Fig. 3.**
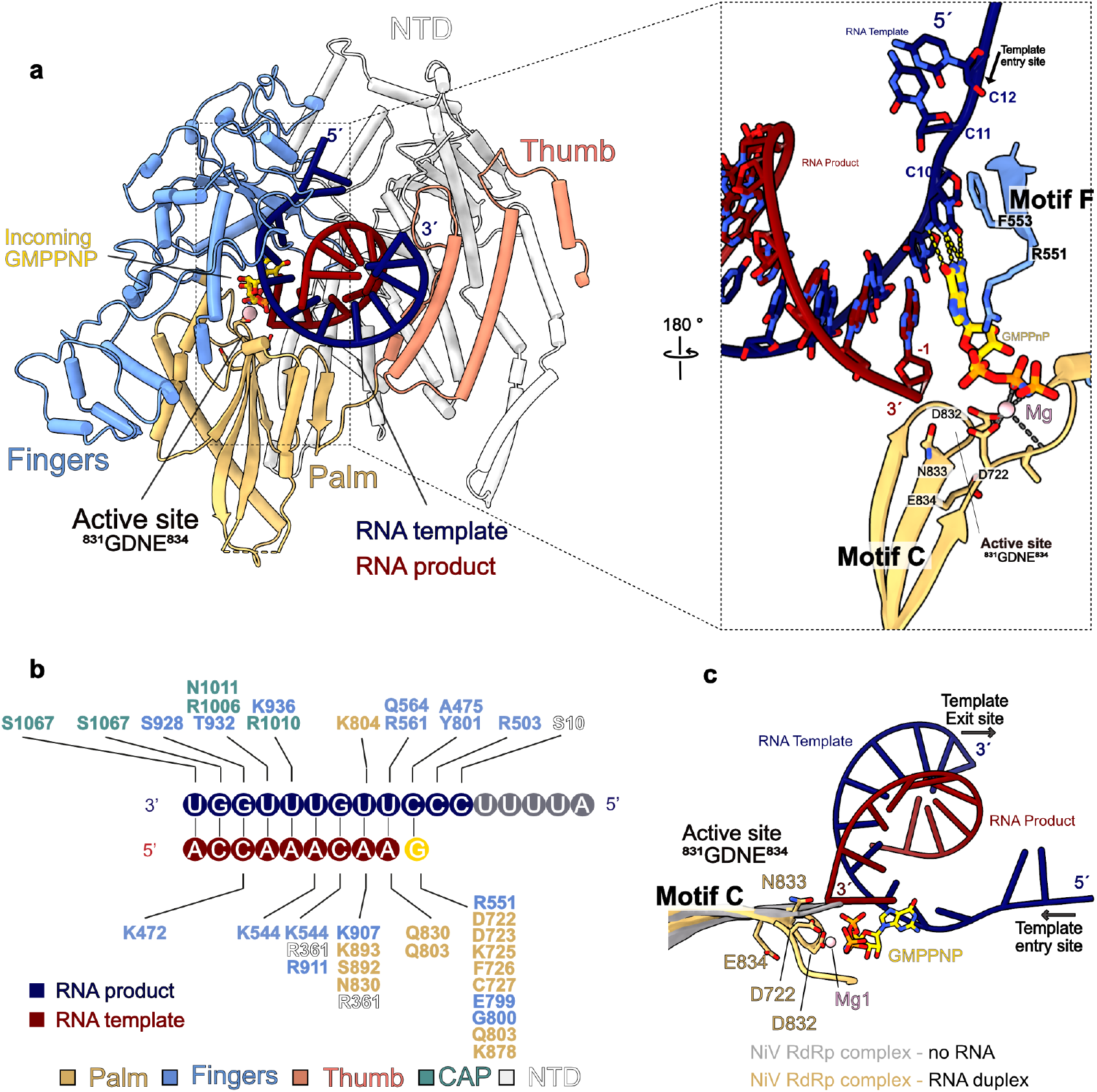
Details of interactions between the NiV RdRp complex and RNA. **a)** Overall fold of the RdRp domain with the fingers, palm, and thumb subdomains colored and labeled as indicated. A close-up view of the active site shows the interactions between the RNA duplex and incoming NTP with motifs C and F. **b)** Schematic of protein-RNA interactions. **c)** Superposition of motif C in the NiV apo- and replicating NiV RdRp structure reveals a conformational change in the active site to accomodate the RNA.

Taken together, the structure of replicating NiV RdRp reveals how it interacts with template and product strands a as well as incoming NTPs during early RNA synthesis and shows that the architecture of the active site is highly conserved.

### Structure of the CD-MTase-CTD module of NiV RdRp

The cryo-EM reconstruction of actively replicating NiV RdRp also reveals the structure of the NiV L CD, MTase and C-terminal domains, which are flexible in the apo state of the enzyme but become ordered in the RNA-bound state (**Figure 2, Figure 4a**). The CD features a conserved compact architecture composed mainly of alpha helices (**Figure 4a, Supplementary Figure 7a**). It is connected to the adjacent CAP and MTase domains by linkers at both ends of the CD (residues 1452-1469 and 1746-1758). The N-terminal linker is partially ordered (res. 1452-1463) and the C-terminal linker is disordered, suggesting that they may act as flexible hinges between the domains, as previously suggested^14^. The MTase domain of NiV adopts a characteristic fold comprised of an eight-stranded β-sheet flanked by helices, which is structurally conserved across nsNSV RdRps (**Supplementary Figure 7b**). It serves as a dual-function enzyme during co-transcriptional capping by methylating the GTP cap first at the 2′-O position and then at the N7 position^26,30,31^. The binding site for the methyl donor S-adenosylmethionine (SAM) contains a conserved GxGxG motif (residues 1843-1847), and a second set of conserved charged residues K-D-K-E (K1821, D1940, K1976, E2013) forms the catalytic tetrad required for methyl group addition (**Figure 4b, Supplementary Figure 4**)^31–33^. The CTD domain, while not strongly conserved at the sequence level, is structurally homologous to those found in other nsNSV RdRps (**Supplementary Figure 4, 7b**). Its most C-terminal helix stacks against the MTase domain, forming a narrow, positively charged RNA-binding groove. It contains a partially conserved KKG motif (K2232, K2236, G2239) which faces towards the active site of the MTase domain and has been implicated in both methylation and capping (**Figure 4b**)^31,34^. Thus, the CTD likely plays in important role in capping during NiV transcription, as reported for other nsNSVs^31,34–36^.

**Fig. 4.**
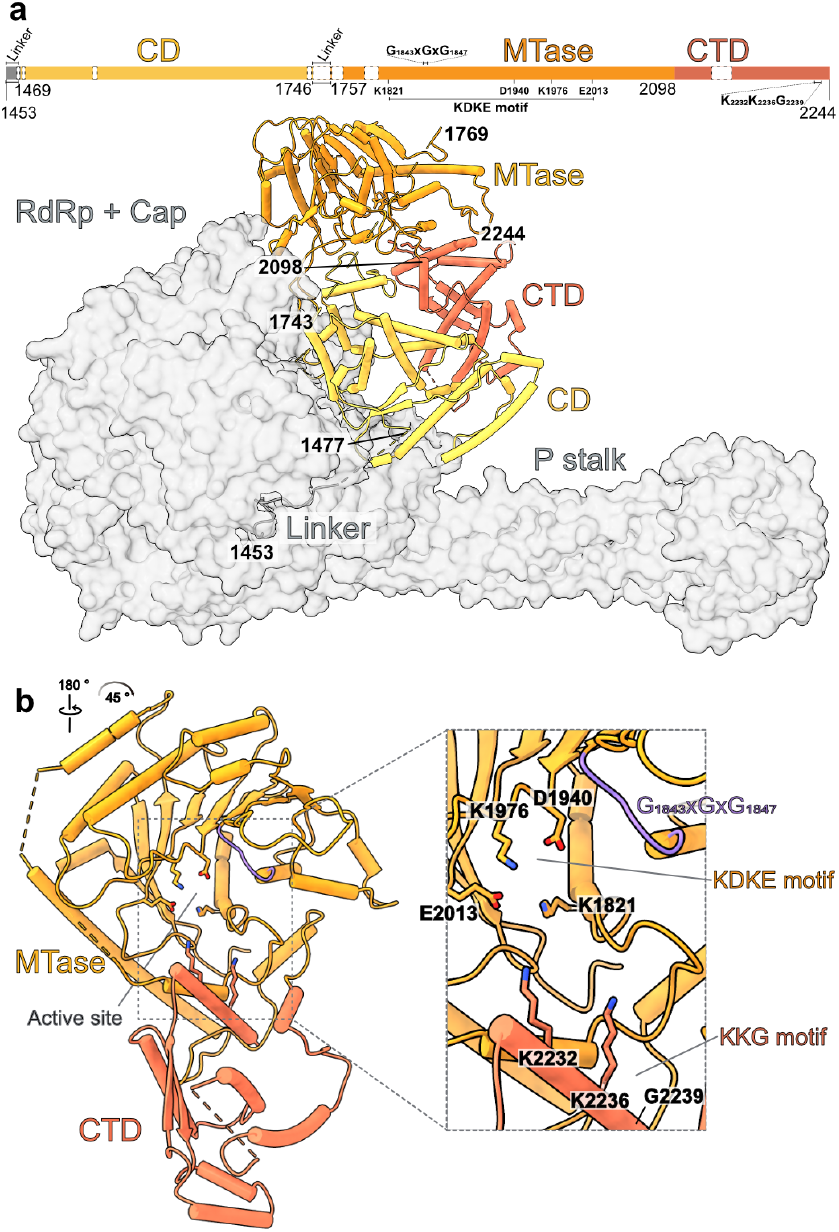
Structure of the NiV L C-terminal domains. **a)** Schematic domain representation of the NiV L C-terminal domains with key catalytic residues indicated. Unmodelled regions are indicated as white dashed boxes. The CD, MTase and CTD domain are shown as cartoon while the RdRp, CAP and P-stalk are shown as transparent gray surface. **b)** MTase-CTD domain with a close-up view of the MTase active site. The KDKE and KKG motifs are shown as sticks, and the GxGxG motif is highlighted in purple.

In previously reported structures of apo nsNSV RdRp complexes, the CD, MTase and CTD were either not visible or observed to adopt different orientations relative to the RdRp and CAP domains^6^. Comparisons to these structures shows that the arrangement of the C-terminal part in the RNA-bound NiV RdRp complex most closely resembles that seen in the hPIV3 RdRp complex^14^, while differing from the orientation observed in the PIV5 RdRp complex (**Supplementary Figure 7c**)^15^. The former was suggested to represent a replication-competent conformation, while the latter was suggested to represent a transcription-competent arrangement due to the closer proximity between the MTase and PRNTase active sites. Thus, the arrangement observed in the RNA-bound NiV RdRp is consistent with an early replicating complex.

In summary, the structure of RNA-bound NiV RdRp reveals the architecture and orientation of the flexible CD-MTase-CTD module during early elongation.

### Conformational rearrangements upon RNA binding and synthesis

Comparison between our two structures reveals conformational rearrangements that occur during the transition from the apo to the RNA-bound state of NiV RdRp. In the RNA-bound state, the priming loop (residues 1266-1290) and intrusion loop (1337-1362) become partially ordered (**Figure 5a**). Both loops run along the interface between the RdRp and CAP domains and mediate interactions between the two as well as the CD (**Figure 5b**). In addition, both loops traverse the active site cavity of the RdRp domain and are located close to the upstream edge of the RNA duplex and the 5’ end of the nascent RNA. The intrusion loop remains extruded from the RdRp active site, thus leaving space for the RNA product and template to fit in. Intriguingly, in both loops the parts in direct proximity to the putative RNA exit channel (res. 1274-1281 of priming loop and 1340-1355 of intrusion loop) show weak density, indicating that they remain flexible. This includes the HR motif in the intrusion loop, which interacts with the 5’ end of the RNA during capping^27^. While this results in a seemingly unobstructed tunnel for the RNA to exit towards the solvent, it is possible that the disordered regions occupy this path. Thus, further elongation of the RNA may trigger rearrangements in the enzyme. Additional structural changes occur in the RdRp domain of NiV L. In particular, the supporting loop becomes fully ordered and adopts a different conformation and the supporting helix, which is mobile in the apo structure, exhibits clear ordered density in the RNA-bound state (**Figure 5c, Supplementary Figure 3b**). These elements interact with the template RNA, the finger domain and with the shifted motif C (**Figure 3c)**. The supporting helix in turn interacts with and may stabilize the priming loop as well as the CD domain (**Figure 5c**).

**Fig. 5.**
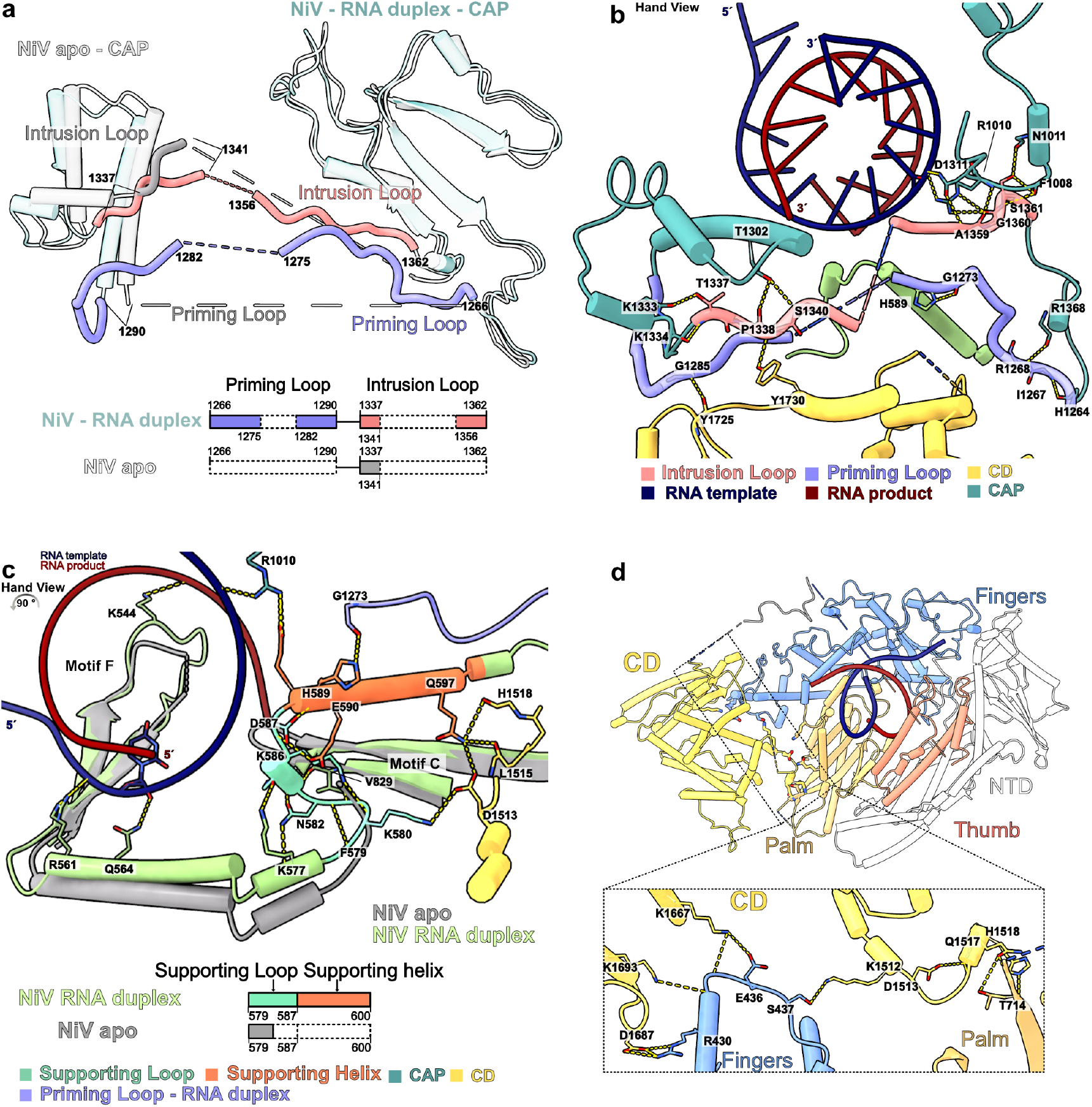
Rearrangements in NiV RdRp upon RNA binding. **a)** Structural and schematic comparison of the priming and intrusion loop in NiV apo and replicating RdRp complex. Both loops become partially ordered in the replicating NiV RdRp complex. **b)** Detailed view of the interactions between the intrusion and priming loops and other elements in the replicating NiV RdRp complex. Key interacting side and main chains are depicted as sticks, with hydrogen bonds highlighted by yellow dashed lines. **c)** Structural overlay and schematic comparison of the supporting helix and loop in the NiV apo (grey) and replicating RdRp complex (colored). The supporting loop and helix become ordered in replicating NiV RdRp complex. Interactions involving the supporting helix and -loop are shown as yellow dashed lines. **d)** Cartoon representation of the RdRp and CD domains in the replicating NiV RdRp complex, with a close-up view of the interactions between them. Hydrogen bonds are depicted as yellow dashed lines, and interacting residues are shown as sticks.

These rearrangements in NiV L may explain how RNA-binding leads to ordering of the flexible C-terminal domains. Several structural elements that undergo conformational changes or become stabilized upon RNA binding interact with the CD domain, in particular the intrusion loop, priming loop, fingers domain, supporting loop and helix, and CAP domain (**Figure 5b,c,d**). These interactions likely stabilize the CD, which in turn may lead to ordering of the MTase and CTD in the observed orientation. Consistent with this, other nsNSV RdRp structures with ordered C-terminal domains also show ordered priming and intrusion loops as well as supporting loop and helices, and similar conformations of motif C (**Supplementary Figure 7d**).

Taken together, our data reveal structural rearrangements in the RdRp core upon RNA binding and early elongation and suggest how these rearrangements may trigger large-scale changes in the architecture of the enzyme.

### Model for NiV RNA synthesis

The structures of the apo and replicating NiV RdRp complex allow us to propose a structural model for the early steps of Nipah virus genome replication. First, the RdRp is in its apo state, with disordered C-terminal domains as well as priming and intrusion loops and supporting loop and helix (**Figure 6a**). Next, it binds to the leader RNA to form an initiation complex (**Figure 6b**). Structural modeling based on the recently reported EBOV^17^ and RSV^18^ initiation complexes shows that no major structural rearrangements are necessary for this, as the template strand can be accommodated in the active site without clashes (**Supplementary Figure 6f**). Whether or not the priming loop transiently occupies the RdRp active site to facility initiation remains unclear. After initiation, the complex transitions into an early elongation state (**Figure 6c**). In this state, the supporting loop and helix are fully ordered, and the priming loop and intrusion loop are partially ordered, which leads to stable association of the previously mobile C-terminal domains. Whether these ordering events occur already during initiation or only after the synthesis of several nucleotides of product RNA is not clear. In this state, the complex can synthesize at least 9 nt of product RNA. Whether further elongation of the RNA then triggers additional conformational changes in the complex and how the ensuing events differ between replication and transcription remains to be determined.

**Fig. 6.**
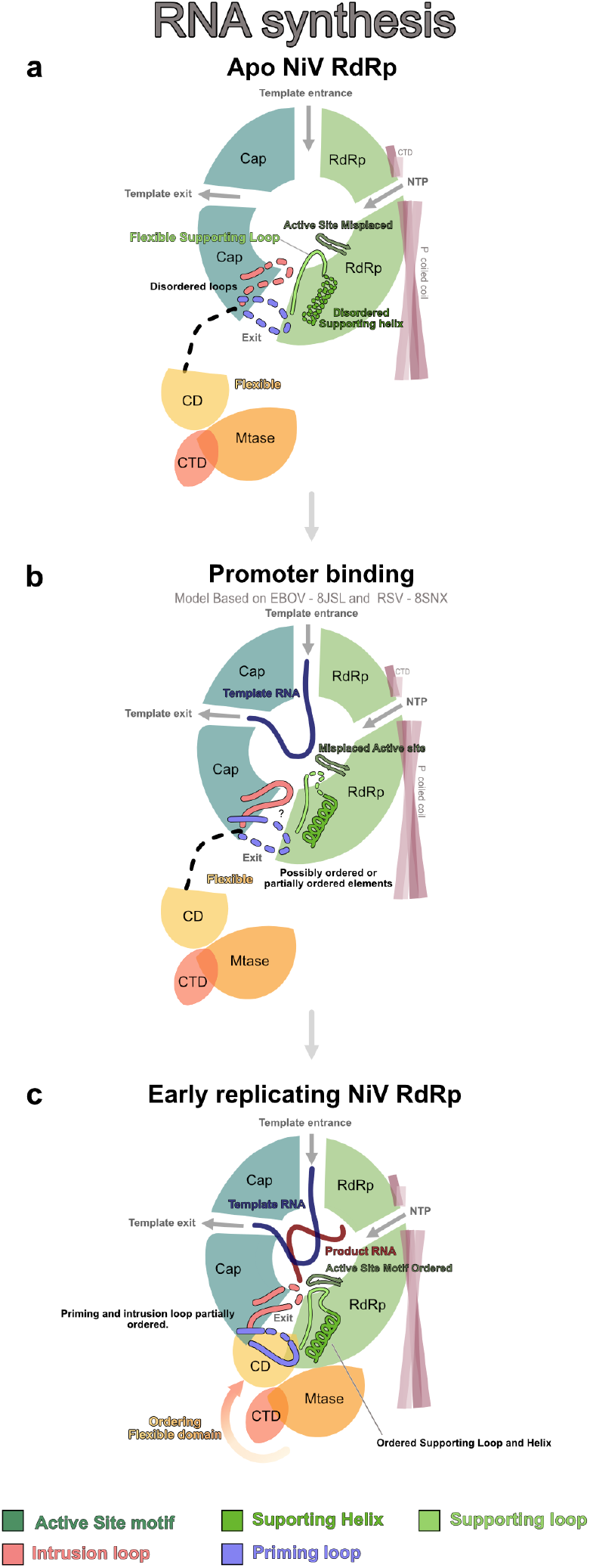
Structure-based model for early steps of NiV replication. **a)** In the **apo state**, the active site motif C is displaced and the intrusion, priming loop, and supporting helix are disordered. The C-terminal domain is flexible. **b)** Model of the **promoter bound state**. The model is based on structures of Ebola virus (EBOV)^17^ and Respiratory Syncytial Virus (RSV)^18^ RdRps in complex with promoter RNA (PDB IDs: 8JSL and 8SNX, respectively). No major rearrangements are required to accomodate the template RNA. The supporting loop, supporting helix, and priming loop may become partially or transiently stabilized. **c)** In the **early replicating state**, motif C shifts to accommodate the RNA product and the supporting helix and loop become fully ordered. The priming and intrusion loops become partially ordered. The supporting loop and helix interact with motif C, the CD, the CAP domain, and the intrusion loop. The C-terminal domains becomes stabilized and ordered.

## DISCUSSION

Here, we present structures of the NiV RdRp complex in its apo and actively replicating state. These structures not only provide the first molecular insights into the Nipah virus replication machinery, but also represent the first structural snapshots of an actively elongating nsNSV RdRp complex.

The apo structure reveals the architecture of the replication machinery of Nipah viruses. It shows that NiV L and four copies of NiV P associate to form a complex which adopts an overall structure that is highly conserved among nsNSV RdRps. Its core is formed by the L protein from which a rod-like stalk formed by the OD domains of four copies of P protrudes. The P-stalk of NiV RdRp exhibits a unique mushroom-like tip formed by helical regions that fold back onto the OD domains, which was also observed in a previous crystal structure of NiV P^8^. Our structure confirms that it also forms in the context of the intact RdRp complex. While the RdRp and CAP domains of L are clearly resolved, the C-terminal CD, MTase and CTD are invisible in the apo structure of the NiV RdRp complex, suggesting conformational flexibility. This is consistent with previously reported apo structures of RSV, HMPV and EBOV RdRps, in which the C-terminal domains were also not visible^11–13^. By contrast, in the apo structures of VSV, NDV, hPIV3 and PIV5 RdRps, these domains were observed to be ordered, but adopt varying orientations with regards to the core^10,14,15,19^. Based on these structures, it has been suggested that the conformational state of the C-terminal domains correlates with distinct functional states of the RdRp complex and is coupled to the conformation of the priming and intrusion loops^6,14^. In particular, in many structures with ordered C-terminal domains, either the priming or intrusion loops invade the active site cavity and would likely clash with an RNA duplex^9,10,14,15,20^. By contrast, in enzymes with disordered C-terminal domains, such as RSV, HMPV or EBOV RdRp, the priming and intrusion loops are either retracted to the CAP domain or disordered, and the active site is unobscured. Consistent with this, in the apo NiV RdRp structure both the priming loop and the intrusion loop are disordered and the C-terminal domains are flexible. During the preparation of this manuscript, two preprints were published that report structures of the apo NiV RdRp complex that are virtually identical to the one presented here^37,38^. In both cases, the CD, MTase and CTD are also not visible, independently confirming that these domains are flexible in the apo state of the enzyme.

The structure of RNA-bound NiV RdRp represents the first structure of a nsNSV RdRp in the actively replicating state. By using a GTP-analog with a non-hydrolyzable bond between the ß and y phosphate groups, we were able to trap an early elongation complex after in situ synthesis of 9 nt of product RNA and with the incoming substrate NTP bound in the +1 site. The structure shows how the enzyme interacts with template and product RNA during early elongation and reveals key residues involved in substrate binding and catalysis. Structural comparisons show that the overall organization of the active site during RNA synthesis is highly conserved across RNA viruses. Our structure of actively replicating NiV RdRp thus provides an important puzzle piece for the mechanistic understanding of nsNSVs and a framework for studying the mechanism of action of nucleoside analog inhibitors.

Comparison between the two NiV RdRp structures reported here reveals the structural rearrangements that accompany the transition from the apo to the RNA-bound state of the enzyme. Upon RNA binding and early elongation, the CD, MTase and CTD become ordered. This is accompanied by rearrangements in the RdRp domain (supporting loop and helix) as well as the CAP domain (priming and intrusion loops). In the apo state of NiV RdRp, the supporting loop and helix as well as the priming and intrusion loops are partially ordered or disordered. In the replicating NiV RdRp complex, the supporting loop and helix become fully ordered and are stabilized by interactions with the RNA and the RdRp domain was well as with the intrusion loop and CD. This is reminiscent of a previous structure of EBOV RdRp incubated with RNA, in which the supporting helix was also shown to be stabilized by interaction with motif C but no continuous density was observed for the supporting loop^11^. In the replicating NiV RdRp complex, the priming and intrusion loops further become partially ordered and run along the interface between the RdRp and CAP domains, near the RNA exit tunnel. Both elements interact with the CD domain and, in conjunction with the supporting loop and helix, likely support its ordering, suggesting a mechanism how RNA-binding in the RdRp core could lead to ordering of the flexible C-terminal domains. Thus, the structure of the replicating NiV RdRp complex reveals the architecture of its previously mobile C-terminal domains and suggests how structural rearrangements during early elongation lead to their ordering.

In summary, our data reveal the structural basis of Nipah virus replication and provide mechanistic insights into nsNSV replication in general. This provides a framework for understanding the mechanisms of replication and transcription of a broad range of human pathogens and may aid in the development of targeted anti-viral therapies.

## METHODS

No statistical methods were used to predetermine sample size. The experiments were not randomized, and the investigators were not blinded to allocation during experiments and outcome assessment.

### Cloning and protein expression

Sequences encoding NiV L and NiV P (NCBI Reference Sequence NC_002728.1) were obtained as codon-optimized variants for expression in insect cells. The sequence encoding NiV L was cloned into the 438-C vector (kind gift from Scott Gradia; Addgene plasmid # 55220; http://n2t.net/addgene:55220; RRID:Addgene_ 55220^44^) in frame with an N-terminal 6xHis tag followed by an maltose binding protein and a TEV cleavage site. The sequence encoding NiV P was cloned into the 438-A vector (kind gift from Scott Gradia; Addgene plasmid # 55218; http://n2t.net/addgene:55218; RRID:Addgene_ 55218^44^) encoding a tagless variant. These vectors were used for subcloning a co-expression vector encoding the 6xHis-MBP-NiV-L and NiV-P proteins. Proteins were expressed in an insect cell expression system using Hi-Five cells as previously described^45,46^.

### Protein purification

After growth, cells were harvested by centrifugation and resuspended in Buffer A (50 mM HEPES pH 8.0, 400 mM NaCl, 6 mM MgCl2, 10% glycerol, 5 mM DTT, and 0.01% Tween 20), supplemented with cOmplete™, EDTA-free Protease Inhibitor Cocktail (Sigma-Aldrich). Cell lysis was performed by sonication, followed by centrifugation at 25,000 rcf for 1 h at 4°C. The soluble fraction was collected and subjected to ultracentrifugation at 235,000 rcf for 60 min at 4°C. The supernatant was filtered through 0.8 µm membranes and applied to a HisTrap HP 5 mL column (GE Healthcare) pre-equilibrated with Buffer A. The column was washed with 10 column volumes (CV) of Buffer A, followed by 5 CV of high-salt buffer (Buffer A with additional 600 mM NaCl). The protein was eluted using Buffer A containing 500 mM Imidazole. For further purification, the eluate was incubated with amylose beads for 2 hours at 4°C. The resin was loaded onto an empty column and washed with 10 CV of Buffer A to remove unbound proteins. The protein of interest was eluted with Buffer A containing 200 mM maltose. The eluted protein was digested with 1 mg of His-tagged TEV (12 h at 4°C). The digested protein was reapplied to a HisTrap column equilibrated with Buffer A. The flow-through was diluted to a final NaCl concentration of 150 mM (Buffer B: 50 mM HEPES pH 8.0, 150 mM NaCl, 6 mM MgCl2, 10% glycerol, 5 mM DTT, and 0.01% Tween 20) and applied to a HiTrap Heparin 5 mL column (GE Healthcare) equilibrated with Buffer B. The protein was eluted with a 300 mM NaCl, using a gradient from 150 to 2000 mM. Peak fractions, as identified by SDS/PAGE, were pooled and concentrated using a MWCO 100,000 Amicon Ultra Centrifugal Filter (Merck). The concentrated sample was applied to a Superose 6 10/300 column equilibrated with Buffer A. Fractions containing Nipah L-P protein were concentrated again using a MWCO 100,000 Amicon Ultra Centrifugal Filter (Merck). The protein with a final concentration of 1.5 uM was freshly used for grids preparation and/or aliquoted and flash-frozen and stored at 80 °C until use. Protein identities were confirmed by mass spectrometry.

### Mass spectrometry

Protein bands were excised from the gel, washed, reduced with dithiothreitol (DTT), alkylated with iodoacetamide and digested with trypsin (sequencing grade, Promega) overnight. The resulting peptides were extracted, dried in a SpeedVac vacuum concentrators (Thermo Scientific) and dissolved in 2% acetonitrile/ 0.05% trifluoroacetic acid (v:v).

Peptides were analyzed by electrospray ionization mass spectrometry in a Thermo Orbitrap Exploris 480 mass spectrometer coupled to an UltiMate3000 ultrahigh performance liquid chromatography system (Thermo Scientific) with an in-house packed C18 reverse-phase column (75 µm ID × 300 mm, Reprosil-Pur 120 C18-AQ, 3 μm, Dr. Maisch). Mass spectrometer was equipped with a Nanospray Flex Ion source and controlled by Thermo Scientific Xcalibur 4.4 software. Data were acquired using a Top30 data-dependent acquisition method. One full MS scan across the 350–1400 m/ z range was acquired at a resolution of 60 000, with an AGC target of 300% and a maximum fill time of 25 ms. Precursors with charge states 2–6 above a 1e4 intensity threshold were then sequentially selected using isolation window of 1.6 m/z, fragmented with nitrogen at a normalized collision energy setting of 28%, and the resulting MS2 spectra recorded at a resolution of 15000, AGC targets of 50% and a maximum fill time of 50 ms. Dynamic exclusion of precursors was set to 22 s.

Proteins were identified with MaxQuant 2.6.1.0 by searching Thermo raw files against a database that included sequences of overexpressed Nipah virus proteins, Trichoplusia ni UniProt reference proteome (release 29-05-2024) and common contaminants observed in MS experiments. Two missed cleavages were allowed. C-carbamidomethyl as well as protein N-terminal acetylation and M-oxidation were set as a fixed and variable modifications, respectively.

### RNA synthesis assay

The RNA template derived from the NiV leader promoter (3’-UGGUUUGUUCCCUUUUA-5’) was purchased from Integrated DNA Technologies. The reaction mixture consisted of 20 μM NiV leader, 1.2 μM recombinant L-P complex, and a buffer containing 50 mM HEPES (pH 8.0), 5 mM DTT, 10% glycerol, and 6 mM MgCl2. Reactions were incubated at 30 °C for 10 min and then initiated by the addition of nucleotides (1 mM ATP, 150 mM fluorescein-labelled CTP, and 1 mM GMPPNP or no GMPPNP) to a final volume of 10 μL, followed by incubation at 30 °C for 12 hours. Control reactions were performed in the absence of NTPs.

The reactions were stopped by adding a buffer containing 7 M urea, 50 mM EDTA (pH 8.0), and 1x TBE buffer. The samples were then treated with 6 U of proteinase K (New England Biolabs) at 37 °C for 30 minutes, denatured at 95 °C for 10 minutes, and analyzed on a 20% polyacrylamide denaturing urea gel. The migration products were visualized using a Typhoon phosphorimager (GE Healthcare, Chicago, IL). The lengths of the RNA products were determined by comparing them to a ladder of synthesized FAM-labelled RNA molecules of known lengths.

### Cryo-EM sample preparation and data collection

For the apo NiV RdRp complex, 0.8 µM of freshly prepared Nipah L-P protein were used for cryo-EM grid preparation. For the replicating NiV RdRp complex, 0.6 µM Nipah L-P protein in Buffer B were mixed with a 15-fold molar excess of RNA scaffold and 1 mM ATP, CTP and GmPPnP, followed by incubation on ice for 2 hours prior to cryo-EM grid preparation. For both samples, 3.5 µL were applied to freshly glow-discharged R 2/1 holey carbon grids (Quantifoil) at 4 °C and 95% humidity. The grids were blotted for 5 seconds with a blot force of 5 using a Vitrobot Mark IV (Thermo Fisher Scientific) before vitrification in liquid ethane.

Cryo-EM data collection was performed with SerialEM 47 using a Titan Krios transmission electron microscope (Thermo Fisher Scientific) operated at 300 keV. Images were acquired in EFTEM mode with a slit width of 20 eV using a GIF quantum energy filter and a K3 direct electron detector (Gatan) at a nominal magnification of 105,000x corresponding to a calibrated pixel size of 0.834 Å/pixel. For the apo NiV RdRp data set, exposures were recorded in counting mode for 1.48 seconds with a dose rate of 23 e-/px/s resulting in a total dose of 48.94 e-/Å2 that was fractionated into 50 movie frames. For the replicating NiV RdRp data set, exposures were recorded in counting mode for 2.14 seconds with a dose rate of 16.9 e-/px/s resulting in a total dose of 52.00 e-/Å2 that was fractionated into 50 movie frames.

### Cryo-EM data processing and analysis

Motion correction, CTF-estimation, particle picking and extraction were performed on the fly using Warp^48^.

An overview of the cryo-EM processing workflow for the apo NiV RdRp dataset is depicted in Supplementary Figure 2. Raw micrographs were split into four batches for initial processing. Particles were extracted with a box size of 480 px and binned threefold. All processing steps were carried out in Cryosparc v.4.5.3^49^ unless stated otherwise. Particles from the first two batches were subjected to initial 2D classification and ‘good’ and ‘junk’ classes were selected manually and each used for ab initio model generation. The resulting best model from the ‘good’ particles and the four ‘junk’ models were then used as input references for a supervised heterogeneous refinement job with all particles. To obtain an improved reference model, particles belonging to the ‘good’ class (∼ 36%; 3.6 million particles) were then again subjected to 2D classification with subsequent ab initio model generation from ‘good’ and ‘junk’ classes. The resulting best model from the ‘good’ particles and the ‘junk’ models were then used as input for a supervised heterogeneous refinement job with the unbinned particles from the good class of the first heterogeneous refinement. Particles belonging to the ‘good’ class (∼ 71 %, 2.5 million particles) were then subjected to unsupervised heterogeneous refinement with 5 classes. Particles belonging to the best class (∼ 22 %, 575 thousand particles) were subjected to non-uniform refinement, which resulted in a reconstruction at a nominal resolution of 2.5 Å, which showed substantial directional anisotropy most likely due to limited view distribution. This map was used as reference together with the models from the first ‘junk’ ab initio job for a supervised heterogeneous refinement using the unbinned particles from the good class of the first supervised heterogeneous refinement as input. Particles belonging to the ‘good’ class (∼ 70 %; 2.5 million particles) were subjected to another round of supervised heterogeneous refinement (4 classes). Particles belonging to the best class (∼24 %; 590 thousand particles) were exported to Relion 5.0^40^ and subjected to 3D refinement with blush regularization^50^ enabled. This resulted in an isotropic reconstruction at an overall resolution of 2.6 Å after post-processing.

An overview of the cryo-EM processing workflow for the RNA-bound NiV RdRp dataset is depicted in Supplementary Figure 5. Raw micrographs were split into four batches for initial processing. Particles were extracted with a box size of 480 px. All processing steps were carried out in Cryosparc v.4.5.3^49^ unless stated otherwise. Particles from the first batch were subjected to initial 2D classification and ‘good’ and ‘junk’ classes were selected manually and each used for ab initio model generation. The resulting best model from the ‘good’ particles and the five ‘junk’ models were then used as input references for supervised heterogeneous refinements carried out for each batch separately (except for batch 1+2, which were processed together). Particles belonging to the ‘good’ class (∼ 34 %) were subjected to another round of supervised heterogeneous refinement with one ‘good’ and five ‘junk’ models and the particles belonging to the ‘good’ class from each batch (∼71 %) were pooled and subjected to unsupervised heterogeneous refinement with six classes. This led to clear separation of particles with and without density for the C-terminal flexible domains of NiV L. The particles belonging to the best class with the C-terminal domains ordered (∼ 35 %, 1 million particles) were exported to Relion 5.0^40^ and subjected to global 3D refinement with blush regularization followed by 3D classification without image alignment, T value of 100, K = 6 and a mask around the NiV L core. Particles belonging to the class with best density for the RNA duplex in the active site were selected (∼33 %; 331 thousand particles) and used for 3D refinement with blush regularization^50^. This led to a consensus reconstruction of the RNA-bound NiV RdRp complex at an overall resolution of 2.8 Å. To further improve the resolution in flexible parts of the NiV RdRp complex, MultiBody refinement using three masks (RdRp core, P-stalk and CTD) was carried out, which led to focused maps of these regions with 2.8, 3.6 and 3.1 Å, respectively and improved the density for the distal parts of the P-stalk substantially. The focused maps were combined into a composite map using the vop maximum command in Chimera^51^.

### Model building and refinement

To generate initial models, the structures of NiV L and P proteins were predicted using Alphafold 3 ^52^. The predicted P protein structure was then partially replaced in the model with the crystal structure of NiV P (PDB ID: 4N5B) ^8^, while retaining the XD domain from the Alphafold model. The overall model was subsequently fitted into the maps as a rigid body using UCSF ChimeraX ^53^ and manually rebuilt using Coot ^54^. Refinement was carried out by real-space refinement in Phenix ^55,56^ as well as by molecular dynamics assisted refinement in ISOLDE ^57^. For the Apo NiV RdRp, final refinement was carried out using the local resolution filtered map. For the replicating NiV RdRp, final refinement was carried out using the composite map followed by a single round of ADP refinement against the consensus map. The resulting models were validated using the MolProbity package within the Phenix suite ^58^. Refinement statistics are shown in **Supplementary Table 1**. Figures were prepared with ChimeraX ^53^.

## Acknowledgements

We thank all members of the Hillen Lab for discussion and Christian Dienemann and Ulrich Steuerwald (MPI-NAT cryo-EM facility) for assistance with cryo-EM data acquisition. H.S.H was supported by the Deutsche Forschungsgemeinschaft under Germany’s Excellence Strategy EXC 2067/1-390729940, FOR2848, SFB1190 and SFB1565 (Project number 469281184, P13).

## Author contributions

F.A.S. designed and carried out all experiments and data analysis unless stated otherwise. K.D. cloned NiV L and P and assisted with insect cell culture. O.D. and H.U. carried out mass-spectrometric analysis. H.S.H. designed and supervised research. F.A.S. and H.S.H interpreted data and wrote the manuscript.

## Data availability

The electron potential reconstructions were deposited with the Electron Microscopy Database (EMDB) under accession codes EMD-51402 (apo NiV RdRp), and for the replicating NiV RdRp: EMD-51403 (composite map), EMD-51723 (Map 1-RdRp core), EMD-51724 (Map-2 – P protein), EMD-51725 (Map 3 – Large protein C-terminal) and EMD-51722 (consensus map). The structure coordinates were deposited to the Protein Data Bank (PDB) under accession codes 9GJT (apo NiV RdRp) and 9GJU (replicating NiV RdRp).

## SUPPLEMENTARY TABLE

**Supplementary Table 1.** Cryo-EM data collection, refinement and validation statistics.

|  | Apo Nipah RdRp complex<br>PDB: 9GJT<br>EMD-51402 | Replicating Nipah RdRp complex<br>PDB: 9GJU<br>EMD-51403 |
| --- | --- | --- |
| **Data collection and processing** |  |  |
| Magnification | 105,000 x | 105,000 x |
| Voltage (kV) | 300 | 300 |
| Electron exposure (e-/Å<sup>2</sup>) | 48.94 | 52.00 |
| Defocus range (μm) | 0.5 – 2.5 | 0.5 – 2.5 |
| Pixel size (Å) | 0.834 | 0.834 |
| Symmetry imposed | C1 | C1 |
| Initial particle images (no.) | 9,886,170 | 10,957,464 |
| Final particle images (no.) | 591,312 | 330,750 |
| Map resolution (Å) | 2.6 | 2.8 |
| FSC threshold | 0.143 | 0.143 |
| Map resolution range (Å) | 2.4 – 5.2 | 2.7 – 5.6 |
| Map sharpening *B* factor (Å<sup>2</sup>) | -93 | -47 |
| **Refinement** |  |  |
| Model resolution (Å) | 2.7 | 2.8 |
| FSC threshold | 0.5 | 0.5 |
| Model composition |  |  |
| Non-hydrogen atoms | 14,017 | 20,948 |
| Protein residues | 1,741 | 2,540 |
| Ligands | 2 x ZN | 2 x ZN, 1 x MG, 1 x GNP |
| *B* factors (Å<sup>2</sup>) |  |  |
| Protein | 81.5 | 54.6 |
| Ligand | 96.8 | 21.7 |
| R.m.s. deviations |  |  |
| Bond lengths (Å) | 0.003 | 0.002 |
| Bond angles (°) | 0.541 | 0.481 |
| Validation |  |  |
| MolProbity score | 1.19 | 1.32 |
| Clashscore | 3.99 | 5.74 |
| Poor rotamers (%) | 0.44 | 1.03 |
| Ramachandran plot |  |  |
| Favored (%) | 98.02 | 98.41 |
| Allowed (%) | 1.92 | 1.48 |
| Disallowed (%) | 0.06 | 0.12 |

## SUPPLEMENTARY FIGURES

**Supplementary Figure 1:**
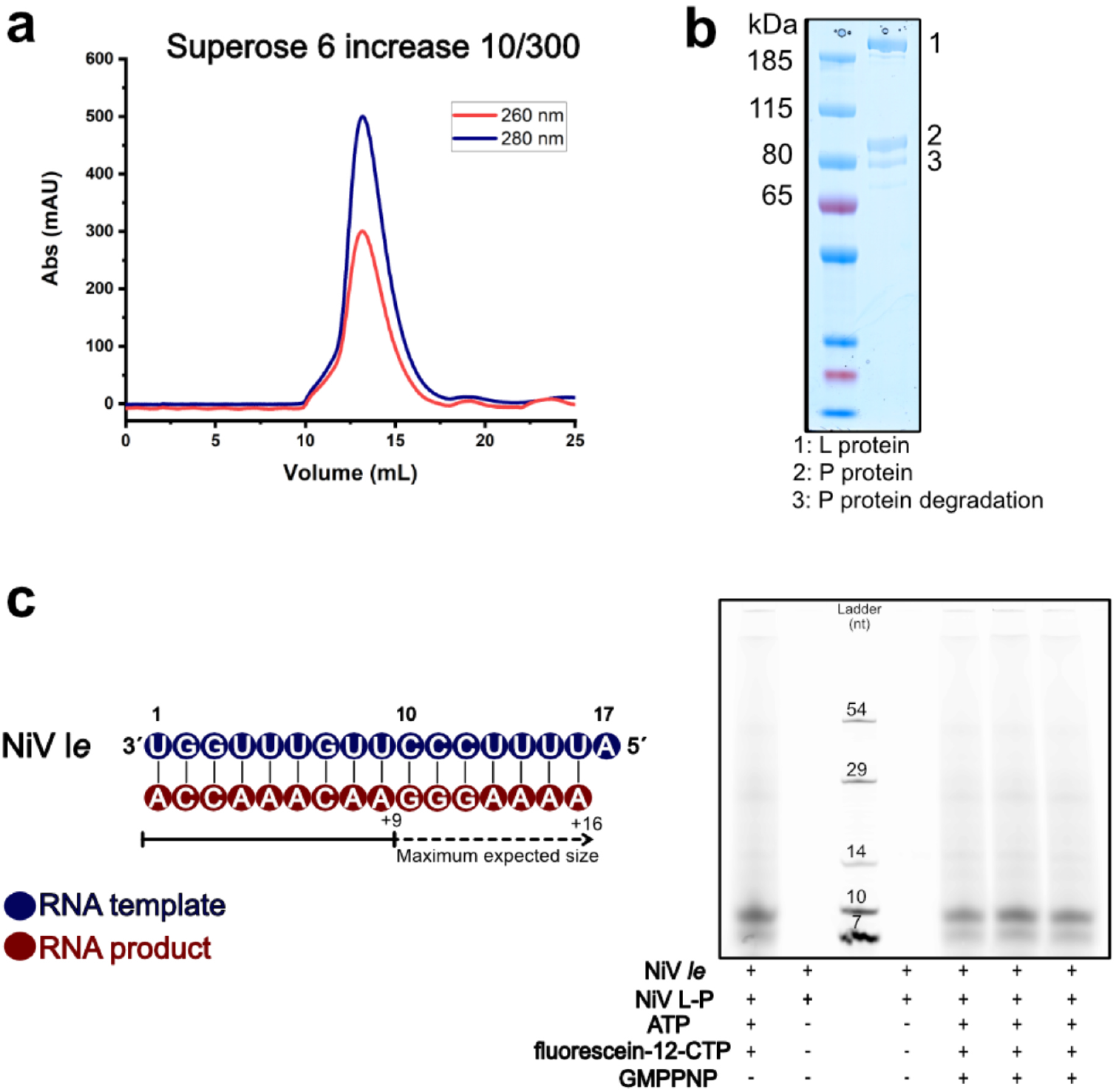
Purification and activity of recombinant NiV RdRp complex. **a)** Size exclusion chromatography profile of the NiV RdRp complex using a Superose 6 Increase 10/300 GL column. The absorbance at 260 nm and 280 nm is shown in red and blue, respectively. **b)** SDS-Page analysis of the purified NiV RdRp copmpex. **c)** RNA synthesis assay using fluorescein-labelled CTP. Products were separated on a 20% urea gel and visualized using with a Typhoon phosphorimager (GE Healthcare, Chicago, IL). RNA product lengths were determined by comparing them to a ladder of synthetic RNA molecules of known sizes.

**Supplementary Figure 2:**
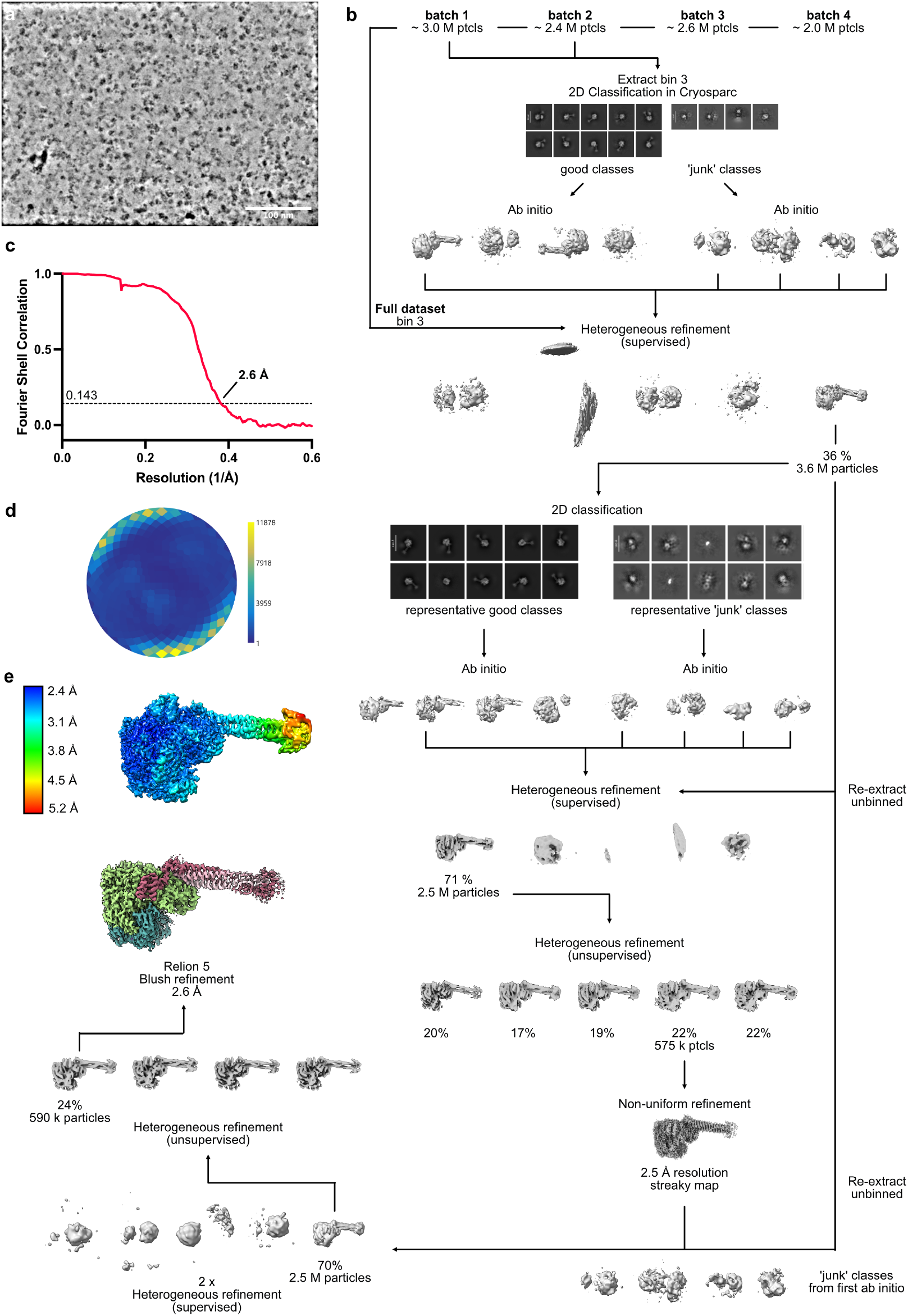
Cryo-EM processing workflow for apo NiV RdRp complex. **a)** Representative denoised micrograph. **b)** Schematic of image processing workflow. See Methods for details. **c)** FSC plot of final reconstruction. **d)** Angular distribution plot created with Warp^39^. **e)** Local resolution filtered map (created in Relion 5.0^40^) colored by local resolution

**Supplementary Figure 3:**
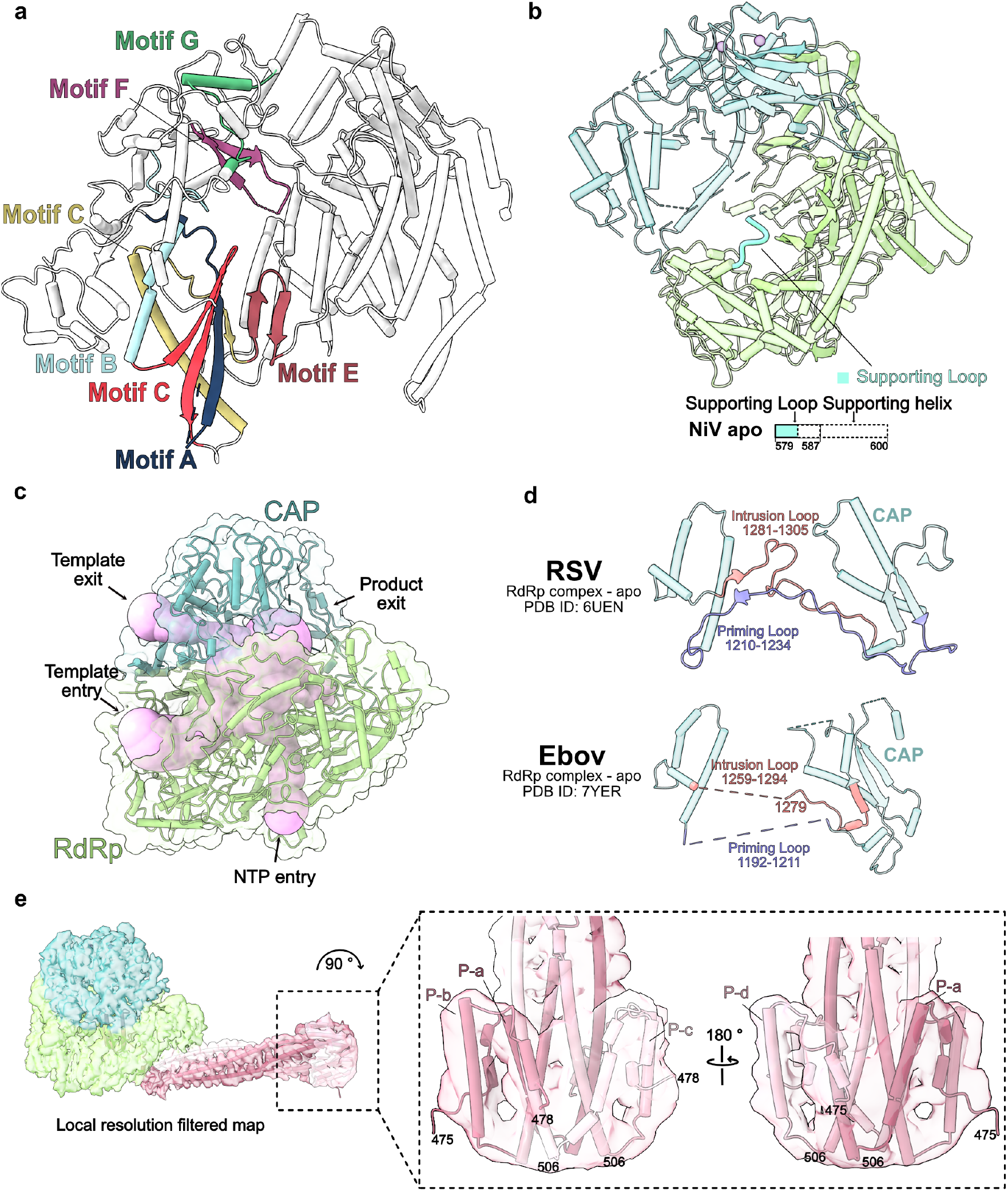
Details and comparison of apo NiV RdRp complex. **a)** Cartoon depiction of conserved catalytic mofifs. The RdRp domain is color-coded with catalytic motifs from A to G displayed and labeled in distinct colors. **b)** Apo NiV L RdRp and CAP domains shown as cartoon with structural and schematic depiction of the supporting helix and loop. **c)** Presumable RNA tunnels within the NiV RdRp complex. Cavities in the complex were calculated using CAVERweb^41^ and are represented by pink spheres. **d)** Comparison of the priming and intrusions loops of Apo NiV RdRp with apo RSV RdRp (PDB ID: 6UEN)^23^ and apo EBOV RdRp (PDB ID: 7YER)^11^. e) Detailed view of the unique P-stalk tip of NiV RdRp. The apo NiV RdRp structure is displayed as cartoon with the local resolution filtered cryo-EM map as transparent surface.

**Supplementary Figure 4:**
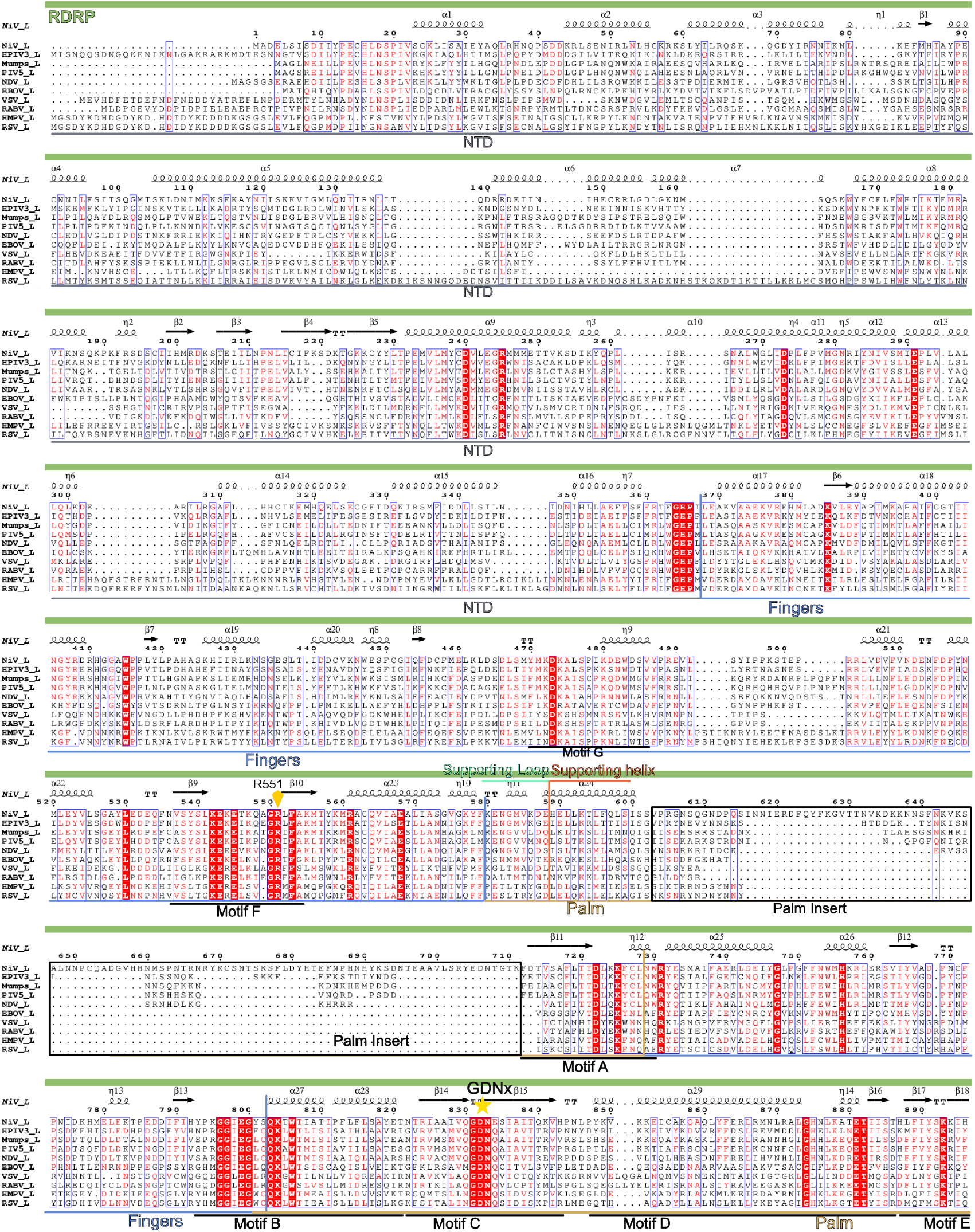
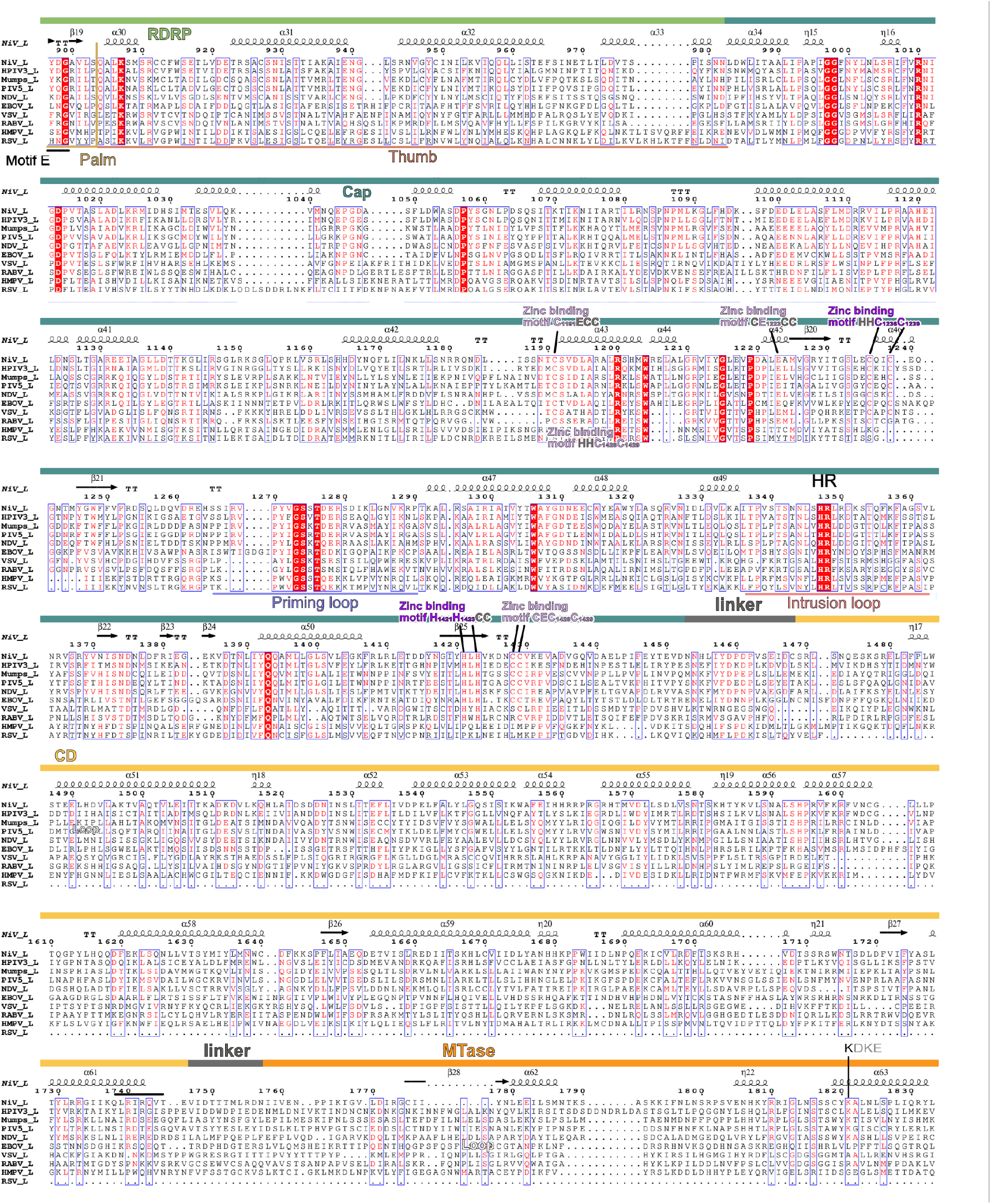
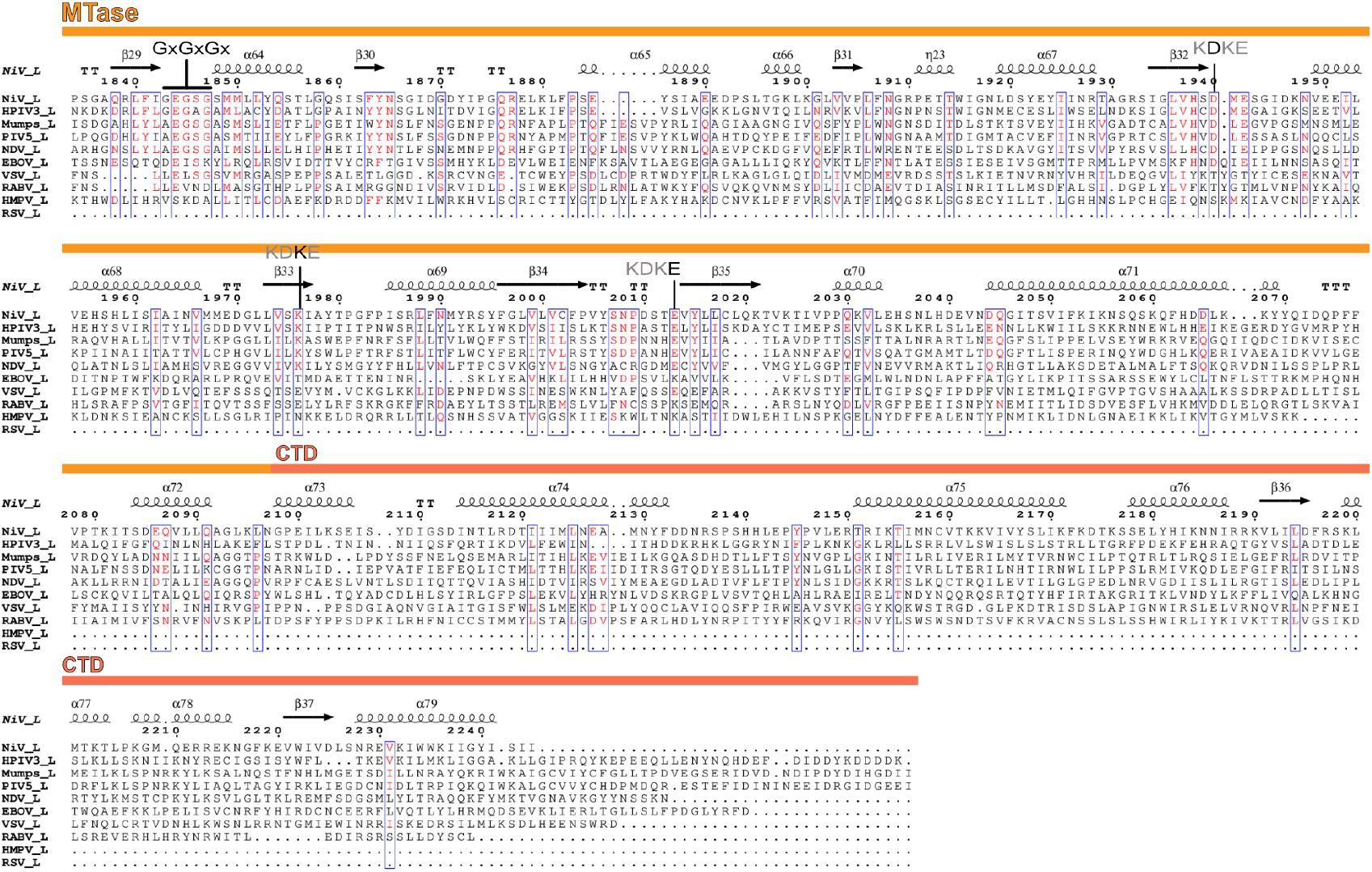
Structure-based sequence alignment of nsNSV L proteins. Sequences for Nipah, hPIV3, Mumps, PIV5, NDV, EBOV, VSV, RABV, HMPV, and RSV. Multiple sequence alignments were performed with MultAlin^42^ combined with secondary structures and alignment results displayed with ESPript3^43^. Domain boundaries, conserved regions, and key motifs are indicated.

**Supplementary Figure 5:**
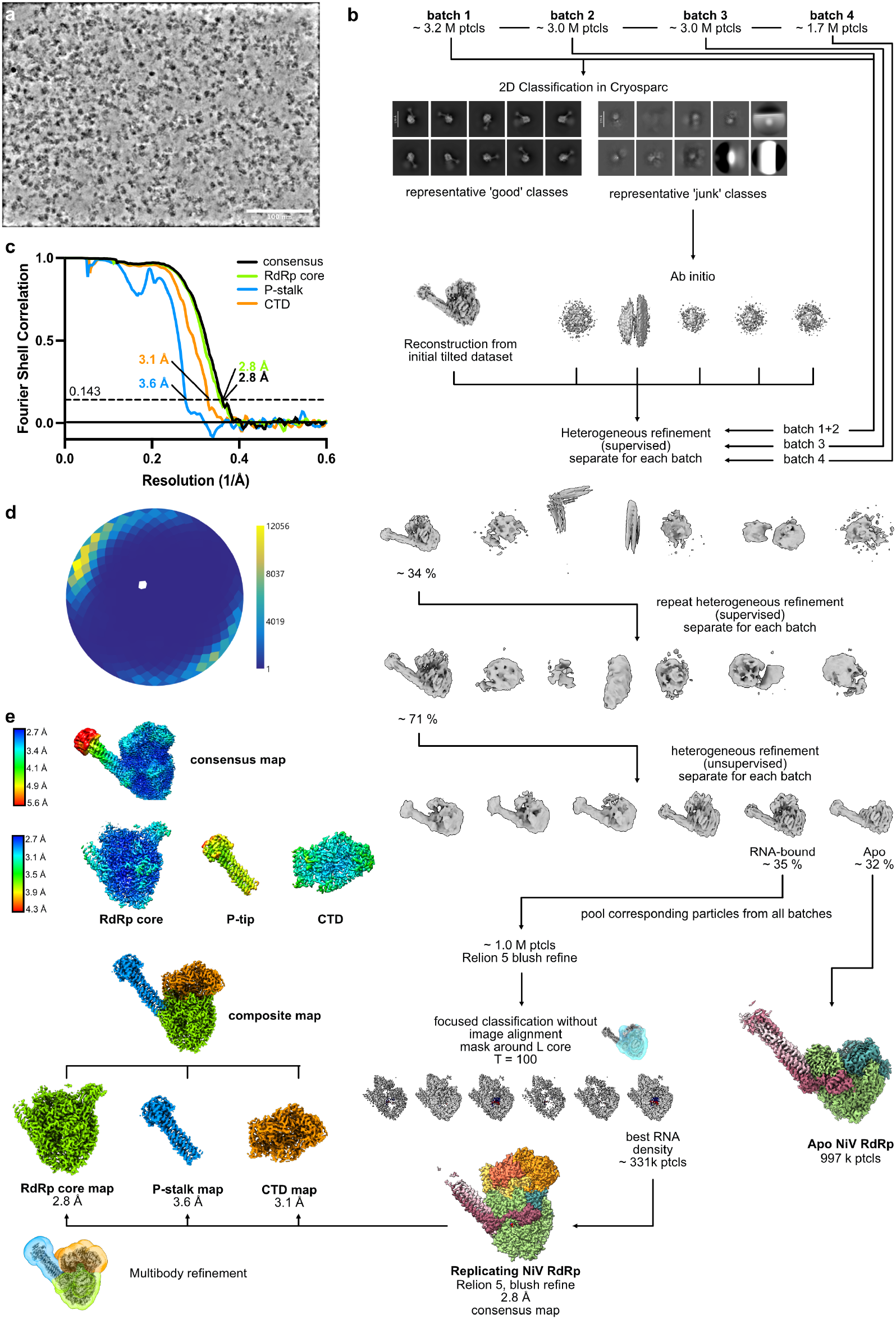
Cryo-EM processing workflow for replicating NiV RdRp complex. **a)** Representative denoised micrograph. **b)** Schematic of image processing workflow. See Methods for details. **c)** FSC plot of final reconstruction. **d)** Angular distribution plot created with Warp^39^. **e)** Local resolution filtered maps (created in Relion 5.0^40^) colored by local resolution.

**Supplementary Figure 6:**
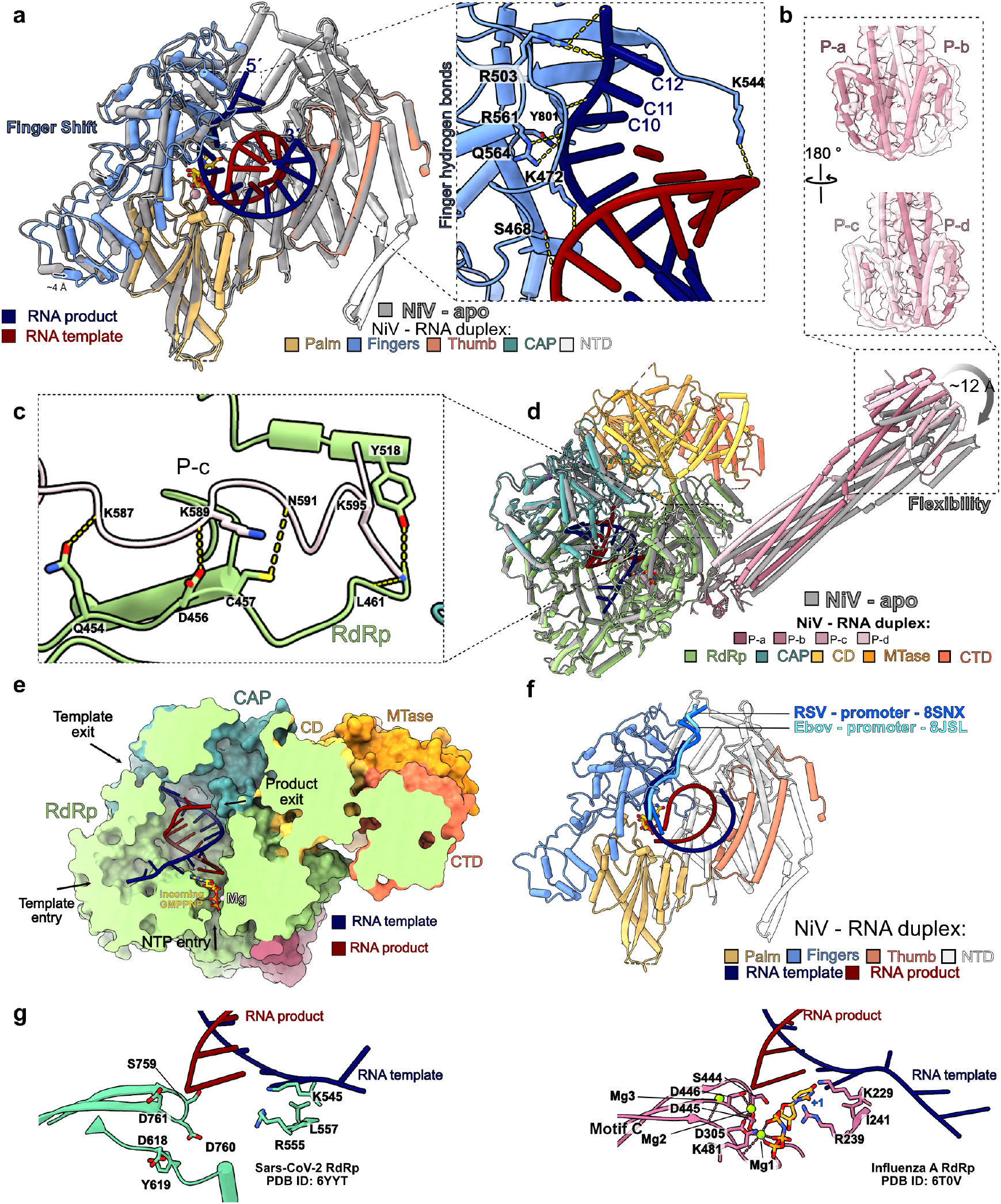
Details and comparison of replicating NiV RdRp complex. **a)**Superimposition of the NiV L RdRp domain in apo (gray) and replicating state (color-coded: finger domain in blue, thumb in salmon, palm in golden yellow, and NTD in gray). The active site and GMPPNP are shown as sticks, with Mg^2+^ ions as pink spheres. The inset displays the interactions between the fingers domain and the RNA template, with hydrogen bonds inidicated by dashed lines. **b)** Close-up view of the P-stalk tip of replicating NiV RdRp. The apo NiV RdRp structure is displayed as cartoon with the composite cryo-EM map as transparent surface. **c)** Close-up view of the interactions between P-c and the L RdRp domain. **d)** Superimposition of apo (gray) and replicating (colored) NiV RdRp complex structures shows conformational flexibility of the P-stalk. **e)** Slice-through view of replicating NiV RdRp complex. The structure is displayed as surface representation with a partially clipped surface. The RNA template and GMPPNP are shown as cartoons and stick, respectively. The template entrance and exit, the NTP entrance and product exit are highlighted. **f)** Comparison to promoter-bound nsNSV RdRp structures. The RdRp domain of replicating NiV L is colored as in panel S6a. The structures of EBOV (PDB ID: 8JSL)^17^ and RSV RdRp (PDB ID: 8SNX)^18^ in complex with promoter RNA were superimposed and the RNA is shown as ribbon. The template RNA follows a similar trajectory in all complexes. **g)** Conserved active site architecture across different RNA viruses. (Left) Close-up view of the SARS-CoV-2 RdRp active site (PDB ID: 6YYT)^28^. (Right) Close-up view of the Bat-Influenza A (PDB ID: 6T0V)^29^ RdRp active site. Residues interacting with RNA are shown as sticks. The incoming NTP is depicted as yellow sticks and Mg^2+^ ion as green spheres.

**Supplementary Figure 7:**
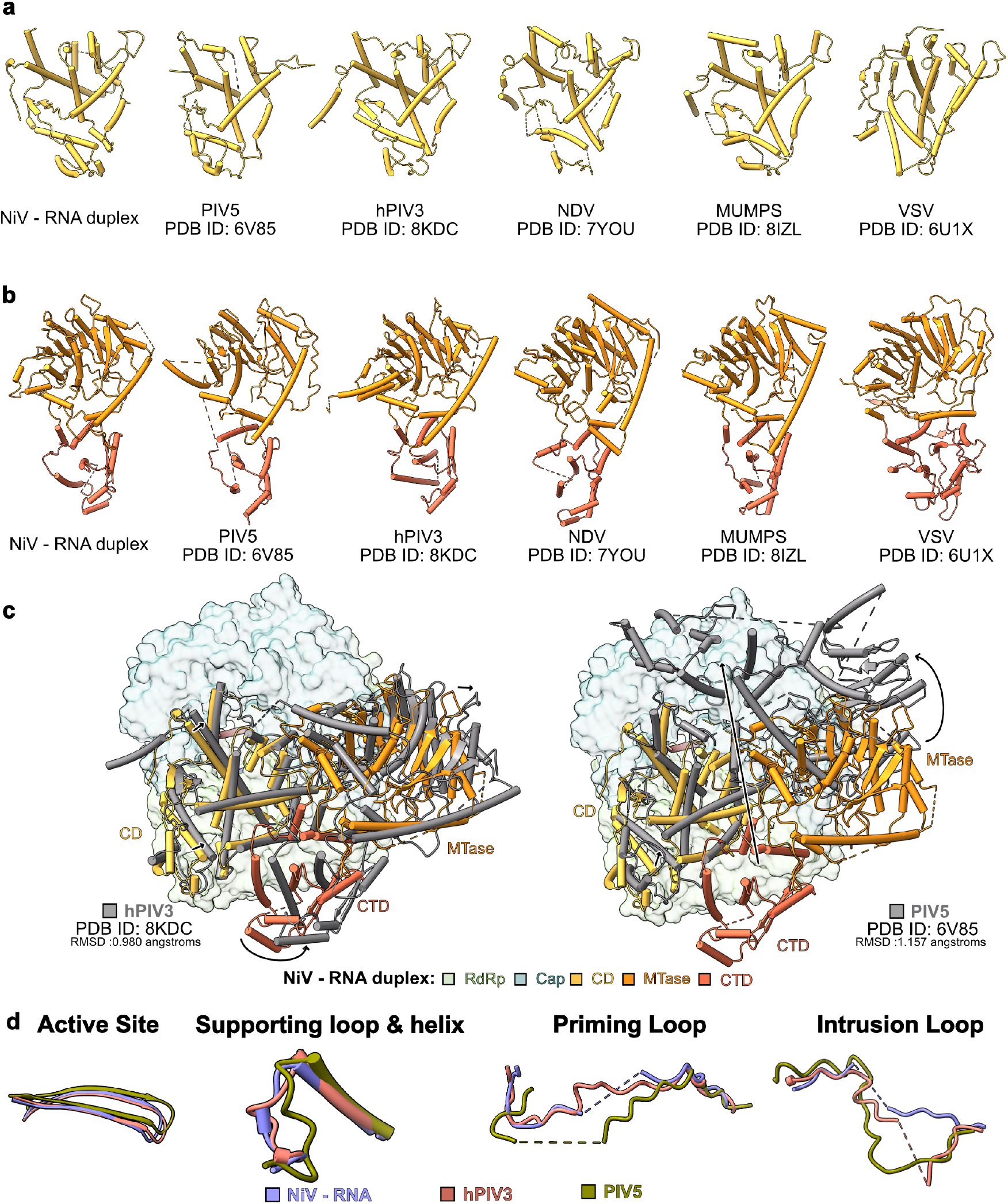
Structural comparison of RdRp complexes from NiV and related viruses. a) Comparison of the CD domains of Nipah virus, parainfluenza virus 5 (PIV5, PDB ID: 6V85)^15^, human parainfluenza virus 3 (hPIV3, PDB ID: 8KDC)^14^, Newcastle disease virus (NDV, PDB ID: 7YOU)^19^, Mumps virus (PDB ID: 8IZL)^16^, and vesicular stomatitis virus (VSV, PDB ID: 6U1X)^20^. (b)Comparison of the methyltransferase (MTase) domain (colored in gold yellow) and CTD (orange) of the viruses described in (a) (c)Superimposition of the L RdRp domain of replicating NiV-RdRp complex with hPIV3 (left, PDB ID: 8KDC, RMSD: 0.980 Å)^14^ and PIV5 (right, PDB ID: 6V85, RMSD: 1.157 Å)^15^. The RdRp and CAP domains are displayed as surfaces in light green and dark cyan, respectively. The CD, MTase, and CTD regions are shown as cartoons, colored as in Fig. 2a for replicating NiV-RdRp and in gray for the other structures. d) Comparison of active site motif C, supporting loop & helix, priming loop, and intrusion loop between NiV, hPIV3, and PIV5 RdRp complexes. Color-coding as indicated.

